# Multiple trans-regulators shape enhancer-promoter hub organization at a multi-enhancer locus

**DOI:** 10.64898/2026.08.19.745721

**Authors:** Sujay Y. Naik, Srijani Roy, Ella Preger-Ben Noon

## Abstract

Developmental genes are frequently regulated by multiple enhancers distributed across large *cis*-regulatory regions. How these enhancers communicate with their target promoter and how their interactions are shaped by distinct developmental transcriptional environments remain incompletely understood. Here, we investigate the chromatin organization of the *Drosophila shavenbaby* locus, a developmental gene controlled by seven distal enhancers. Tissue-specific UMI-4C revealed extensive enhancer-promoter and enhancer-enhancer interactions, including in cell populations where individual enhancers are inactive. Quantitative three-dimensional DNA-FISH revealed compact enhancer-promoter hubs enriched in *shavenbaby*-expressing cells, yet also present in non-expressing cells and prior to expression. Perturbation of *shavenbaby* regulators, transcription factors, and architectural proteins revealed that multiple factors contribute to hub organization. Their relative contributions differed between epidermal populations, indicating that similar hubs can be supported by different combinations of regulators. Perturbations that reduced hub organization were frequently associated with reduced *shavenbaby*-dependent trichome formation. Together, our results identify a robust, multi-factorial enhancer-promoter hub that is shaped by distinct regulatory inputs across developmental contexts.

## Introduction

Developmental genes often rely on the combined activity of multiple enhancers to generate their complete expression pattern. Each enhancer can bind a distinct set of context-specific transcriptional regulators and activate transcription in a defined spatiotemporal pattern^1^, while partial overlap between enhancer activities can confer robustness to gene expression^2–5^. In many cases, these regulatory elements are spread across large genomic regions, far from their target promoters^6,7^. How multiple enhancers regulating the same gene communicate with a common target promoter to coordinate gene expression across different developmental contexts remains poorly understood.

Long-range communication between enhancers and promoters is thought to occur through physical chromatin interactions^7,8^. Over the last decade, chromosome conformation capture (3C)-based studies have revealed widespread interactions between enhancers and promoters across the genome^9–15^. Some of these studies reported multivalent interactions in which several enhancers and promoters engage simultaneously, forming structures referred to as enhancer-promoter hubs or cliques^16–26^. In parallel, imaging studies identified dynamic condensates, referred to as ‘transcriptional hubs’, that are enriched in activators, co-activators, chromatin modulators, and the transcriptional machinery and often coincide with transcriptionally active loci^27–30^.

Whether enhancer-promoter interactions depend on context-specific transcription factor binding and enhancer activity remains debated. Some studies have shown that enhancers and promoters interact preferentially in the tissues and developmental stages in which their target genes are transcribed^31–34^ and may depend on the binding of context-specific transcription factors^14,35,36^. In contrast, other studies documented enhancer-promoter interactions and multivalent enhancer-promoter hubs in cells in which the associated target genes were not expressed, as well as prior to gene activation^12,13,37–41^. More recent studies showed that enhancer-promoter contacts can change during development, with additional enhancer-promoter interactions emerging during differentiation, coinciding with enhancer activation and gene expression^41–43^. These observations raise the possibility that enhancer-promoter interactions are differentially regulated by distinct trans-regulatory environments. However, what determines the organization of multi-enhancer-promoter hubs remains poorly understood. It is unclear which trans-regulators contribute to enhancer-promoter hub organization and whether their relative contributions differ across developmental contexts.

The *Drosophila shavenbaby (svb)* locus provides an ideal system to address these questions in a developmental multi-enhancer locus^44^. *svb* encodes a transcription factor required for embryonic cuticular trichome formation (Fig. 1a)^45–47^. Its complex embryonic expression is controlled by the combined activity of seven enhancers distributed across a ∼90 kb region upstream of the *svb* promoter (Fig. 1b)^2,48,49^. Each enhancer responds to distinct regulatory inputs and drives expression in discrete, sometimes overlapping domains of the embryonic epidermis (Fig. 1c-i). For example, the *E3* and *7H* enhancers drive expression in the ventral epidermis (Fig. 1h-i) and are regulated by Ultrabithorax (Ubx)^50^, whereas the *E6* enhancer drives expression in the dorsal epidermis (Fig 1f) and is activated by Arrowhead (Awh) and Pannier (Pnr)^51^. The partial overlap between enhancer activities provides robustness to *svb* expression under environmental or genetic variation^2^. Recent studies suggest that this robustness may involve multi-enhancer clustering within transcriptional hubs^52^. Specifically, a *svb* enhancer inserted on a different chromosome colocalizes with the endogenous *svb* gene in transcriptional hubs enriched for Ubx^53^ and can compensate for the loss of endogenous *svb* enhancers under extreme conditions^52^. It remains unknown whether endogenous *svb* enhancers interact with one another and with the promoter, how their organization differs between epidermal domains with distinct enhancer activities, and what trans-regulators contribute to this architecture.

**Fig. 1.**
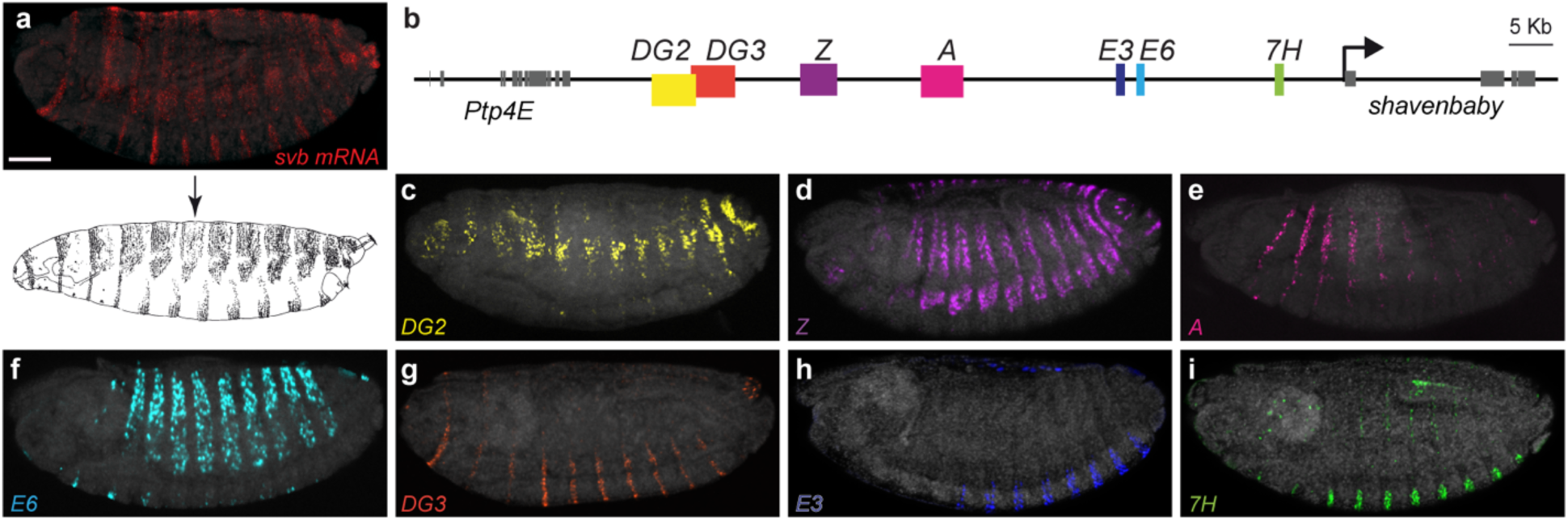
Seven enhancers regulate the embryonic expression of *shavenbaby* (a) Top: Expression pattern of *svb* mRNA in stage 14 *D. melanogaster* embryo visualized by HCR^TM^ RNA fluorescent in situ hybridization. Bottom: Lateral view of a first instar *D. melanogaster* larva. Scale bar, 10 μm. (b) Schematic representation of the *svb* locus showing the positions of the embryonic enhancers (colored boxes). (c–i) Expression of *D. melanogaster DG2::LacZ* (c), *Z::LacZ* (d), *A::LacZ* (e), *E6::LacZ* (f), *DG3::LacZ* (g), *E3::LacZ* (h), and *7H::LacZ* (i) reporter constructs in stage 15 *D. melanogaster* embryos.

Here, we combined tissue-specific chromosome conformation capture assays with quantitative three-dimensional DNA-FISH to determine the organization of the *svb* locus across different trans-regulatory environments in the developing embryo. We found that *svb* enhancers interact extensively with the promoter and with one another, forming multi-enhancer-promoter hubs in which enhancers participate regardless of their activity state. These hubs are present prior to *svb* expression and persist in epidermal cells that do not express *svb*, but occur more frequently in *svb*-expressing nuclei. Perturbation experiments revealed that multiple trans-regulators contribute to hub organization, with distinct effects across developmental contexts. Finally, perturbations that alter hub organization frequently affect *svb*-dependent trichome patterning. Together, our findings suggest that enhancer-promoter hub organization is shaped by the combined contributions of multiple trans-regulators, whose effects vary across developmental contexts.

## Results

### Mapping genomic interactions at the *shavenbaby* locus

Reporter gene assays have shown that *svb* enhancers drive both unique and overlapping expression patterns across multiple developmental contexts^2,48,49,54^. However, how these enhancers interact with the *svb* promoter within their native chromosomal environment remains unclear. To address this, we used cell type-specific chromosome conformation capture with unique molecular identifiers (UMI-4C)^55^ to map chromatin interactions at the *svb* locus in specific epidermal cell subpopulations of *D. melanogaster* embryos.

To isolate nuclei in which individual enhancers are active, we generated two transgenic lines in which dorsal or ventral *svb* enhancer fragments drive nuclear reporters. In the dorsal tester line, the enhancer fragments *E6B*^51^, *Z1.3L*^54^ and *DG2B*^56^ drive *GFP, LacZ,* and *dsRed* expression, respectively, in cell subpopulations with a partial overlap (Fig. 2b). In the ventral line, the enhancers *E3* and *7H* mark complementary ventral epidermal cells (Fig. 2c). Stage 14 embryos from these lines were cross-linked, dissociated, and their nuclei were isolated by fluorescence-activated cell sorting (FACS) based on reporter expression^57^, such that each sorted population corresponds predominantly to nuclei in which a single enhancer is active. These nuclei were then used for UMI-4C^55^ (Fig. 2d).

**Fig. 2.**
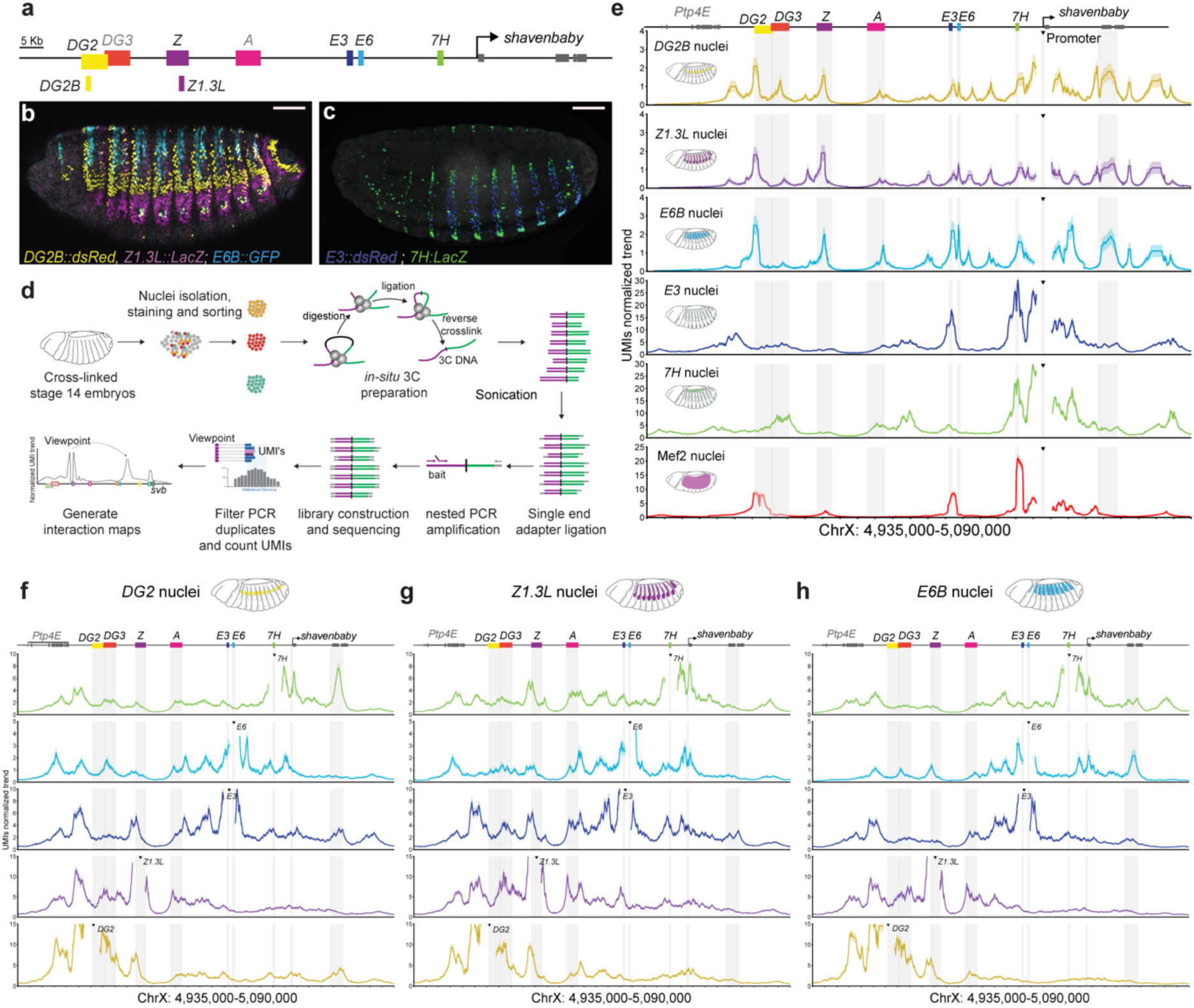
Cell type-specific UMI-4C reveals extensive connectivity across the *shavenbaby* locus (a) Schematic representation of the *svb* locus with the enhancer fragments used to generate the dorsal and ventral tester lines shown below. (b,c) Expression of *DG2B::dsRed* (yellow), *Z1.3L::LacZ* (magenta), and *E6B:: GFP* (cyan) in stage 15 embryos from the dorsal tester line (b), and of *E3::dsRed* (blue) and *7H::LacZ* (green) in stage 15 embryos from the ventral tester line (c). Scale bars, 10 μm. (d) Experimental workflow. Nuclei were isolated from cross-linked stage 14 embryos, sorted by FACS based on reporter expression, and processed for UMI-4C. Libraries were generated following *in situ* restriction enzyme digestion, ligation, sonication, nested PCR amplification, and high-throughput sequencing. (e) Cell type-specific UMI-4C interaction profiles across the *svb* locus using the endogenous *svb* promoter as the viewpoint. Interaction profiles are shown for *DG2B* (yellow), *Z1.3L* (magenta), *E6B* (cyan), *E3* (blue), *7H* (green), and Mef2 (red) nuclei. (f–h) Cell type-specific UMI-4C interaction profiles using the indicated *svb* enhancers as viewpoints in *DG2B* (f), *Z1.3L* (g), and *E6B* (h) nuclei. Profiles are shown for the *7H* (green), *E6* (cyan), *E3* (blue), *Z* (magenta), and *DG2* (yellow) enhancer viewpoints. In (e–h), the y-axis shows UMI-normalized interaction frequencies relative to the total UMI counts. Gray shading indicates the positions of the *svb* promoter and embryonic enhancers, and triangles indicate the corresponding viewpoint in each interaction profile.

We first mapped genomic interactions of the *svb* promoter. In dorsal nuclei, we detected interactions with all dorsal enhancers, as expected (*DG2*, *Z*, *A*, and *E6*; Fig. 2e, Fig. S1a). Notably, the promoter also interacted with ventral enhancers, which are inactive in the sorted subpopulation (Fig. 2e, Fig. S1a). In addition to distal contacts, prominent interactions were detected within the *svb* gene body, including intron-exon junctions. Comparisons of interaction profiles across the three dorsal cell subpopulations revealed both shared and population-specific features. While the overall interaction landscape and promoter contacts with multiple enhancer regions were broadly conserved, interaction profiles differed between subpopulations. In general, interaction profiles showed an enrichment of contacts between the major enhancer in each subpopulation and the promoter (e.g., *E6* in *E6B* nuclei, *Z* in *Z1.3L* nuclei, and *DG2* in *DG2B* nuclei; Fig. 2e, Fig. S1a).

In both ventral cell subpopulations, the *svb* promoter interacted robustly with the ventral enhancers *DG3*, *E3*, and *7H*, while maintaining weaker interactions with some dorsal enhancers (Fig. 2e, Fig. S1a). As in dorsal nuclei, the *svb* promoter also interacted with regions within the gene body. The overall interaction landscape was broadly conserved between *E3*-and *7H-*positive nuclei, with some population-specific contacts. In contrast, non-epidermal *Mef2*-expressing nuclei exhibited reduced interactions with *svb* enhancers (Fig. 2e, Fig. S1a), indicating that extensive enhancer-promoter interactions at the *svb* locus are associated with *svb* expression and epidermal identity.

Next, we examined the chromatin interactions from the perspective of individual enhancers. We generated UMI-4C libraries from sorted dorsal epidermal nuclei subpopulations using viewpoints positioned at the dorsal enhancers *DG2, Z,* and *E6*, and the ventral enhancers *E3* and *7H.* We observed extensive chromatin interactions spanning the upstream regulatory region of the *svb* locus in all dorsal subpopulations examined (Fig. 2f-h, Fig. S1b-d). Each enhancer displayed strong local interactions, together with contacts to the *svb* promoter and to other enhancers. These interactions extended across the locus, indicating that the *svb* enhancers operate within a broader chromatin interaction network rather than as isolated elements. While a common interaction landscape was observed across subpopulations, individual contact patterns differed between cell populations, indicating cell type-specific modulation of enhancer connectivity. Notably, enhancer-enhancer interactions were detected even in subpopulations in which those enhancers are not active (Fig. 2f-h, Fig. S1b-d).

Together, these results demonstrate that the *svb* enhancers and promoter form extensive enhancer-promoter and enhancer-enhancer interactions, suggesting that the *svb* regulatory region is organized as a multi-enhancer enhancer-promoter hub. This organization persists across different regulatory contexts, including in cells where individual enhancers are inactive.

### Spatial organization of the *svb* regulatory region revealed by multicolor DNA-FISH

While UMI-4C revealed extensive interactions between the *svb* promoter and multiple enhancers, it measures pairwise interactions across populations of nuclei and cannot determine whether these interactions occur simultaneously within the same nucleus. To examine the spatial organization of the *svb* regulatory region at single-cell resolution, we performed multicolor DNA-FISH.

We focused on the *svb* promoter (*P*) and the dorsal enhancers *DG2* and *E6*, located approximately 75 kb and 23 kb from the promoter, respectively, providing optimal spacing between the three loci (Fig. 3a). We designed DNA-FISH probes spanning ∼4 kb regions encompassing each element, enabling simultaneous visualization of the promoter and distal enhancers in single nuclei. To ensure that measured distances reflect intra-allelic interactions, analyses were carried out in stage 14 male embryos, which carry a single copy of the X-linked *svb* gene.

**Fig. 3.**
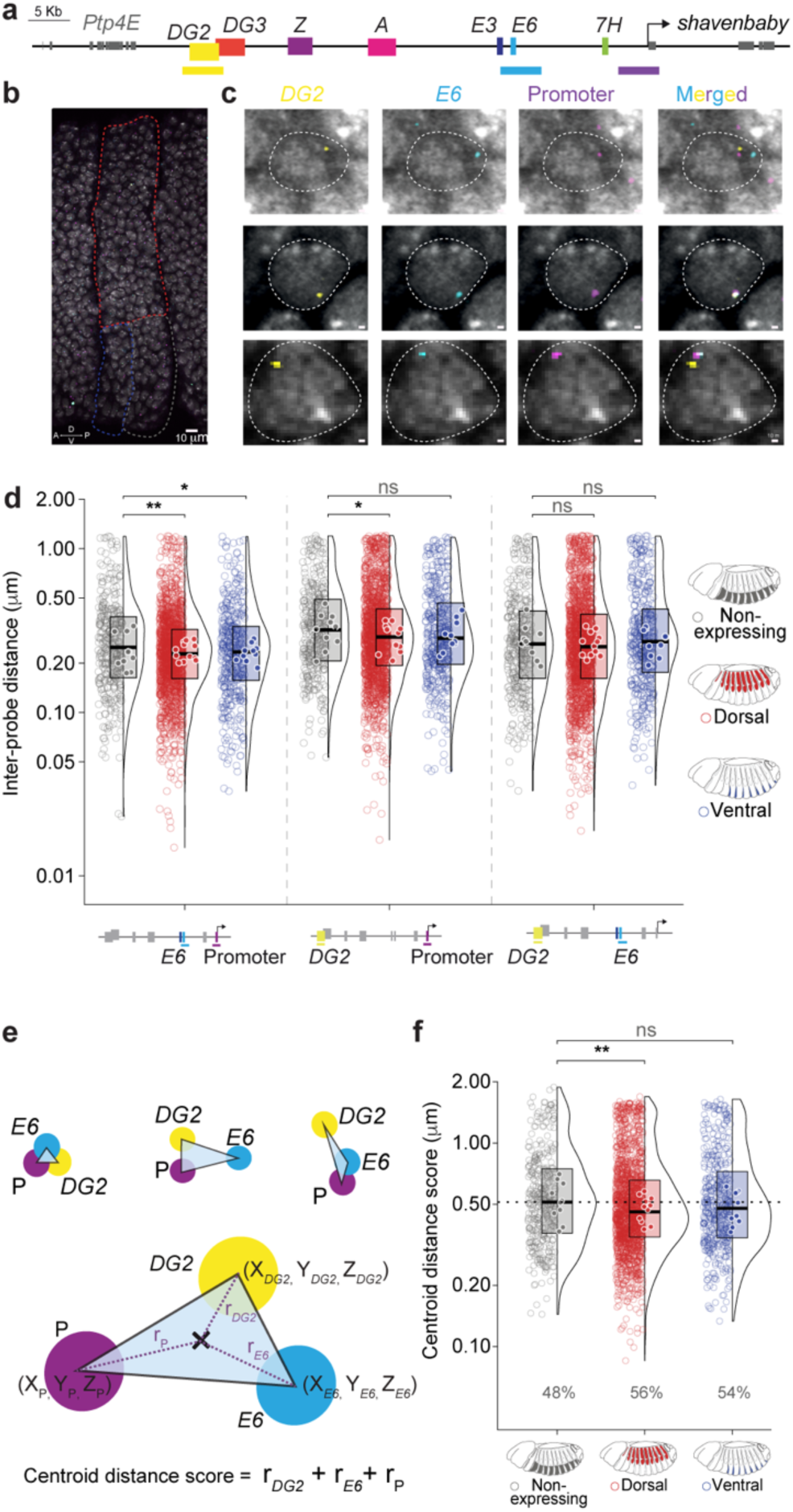
Three-color DNA-FISH reveals a multi-enhancer-promoter hub at the *shavenbaby* locus (a) Schematic representation of the *svb* locus showing the positions of the DNA-FISH probes (color-coded below). (b) Representative DNA-FISH images of abdominal segments A3-A5 of stage 14 male embryo. Non-expressing, dorsal *svb*-expressing and ventral *svb*-expressing regions used for analysis are marked by gray, red and blue dashed lines, respectively. Scale bar – 10 μm (c) Representative DNA-FISH images of single nuclei. Dashed outlines indicate segmented nuclei. Insets show the detected *DG2, E6*, and *svb* promoter signals together with merged images illustrating representative three-locus configurations. Scale bars, 10 μm (d) Pairwise inter-probe distances at the *svb* locus. Three-dimensional pairwise distances between *E6* and the *svb* promoter (left), *DG2* and the *svb* promoter (middle), and *DG2* and *E6* (right) in non-expressing (grey, *n* = 349, *N* = 10), dorsal *svb*-expressing (red, *n* = 1622, *N* = 12), and ventral *svb*-expressing (blue, *n* = 427, *N* = 10) nuclei. For each probe pair, violin plots show the distribution of pairwise distances, box plots indicate medians and interquartile ranges, open circles represent individual nuclei, and solid circles denote embryo-level median values. Statistical significance was assessed using two-sided Wilcoxon rank-sum tests with Benjamini-Hochberg correction (\**P* ≤ 0.05, \*\**P* ≤ 0.01, n.s., not significant). Distances are shown in micrometers (μm). (e) Schematics showing the relative positions of the *DG2* enhancer, *E6* enhancer, and the *svb* promoter (P), the triangle connecting their centroids (top), and the centroid distance score (bottom), calculated as the sum of the distances between the triangle centroid and each locus. (f) Centroid distance scores for the same nuclei shown in (d). Lower values indicate more compact three-locus configurations. Percentages indicate the fraction of nuclei with centroid distance scores below 0.5 μm. Violin plots, box plots, individual nuclei, embryo-level medians, and statistical testing are presented as in (d).

Epidermal nuclei were classified based on their position along the dorso-ventral axis of abdominal segments A3-A7 (Fig. 3b). Dorsal *svb*-expressing nuclei were defined as the ∼25 rows of epidermal cells extending ventrally from the amnioserosa in each segment, whereas ventral *svb*-expressing nuclei were defined as ∼4 columns of nuclei in the ventral epidermis located immediately posterior to the anterior boundary of each segment. The intervening epidermal region was classified as non-expressing (Fig. 3b).

Consistent with the interaction patterns detected by UMI-4C, many epidermal nuclei showed close spatial proximity of the three elements, forming compact enhancer-promoter clusters (Fig. 3c). Representative images also illustrate that the three probe signals can be readily resolved when spatially separated (Fig. 3c). To quantify the spatial organization of the locus, we used complementary geometric measures to assess both pairwise relationships and overall hub organization. We first quantified the three-dimensional pairwise distances between the centroids of the probe signals (Fig. 3d), which showed broad distributions across all cell populations. In non-expressing nuclei, the median pairwise distances between the two enhancer probes and the *svb* promoter ranged from 228 to 319 nm, indicating that these regulatory elements are already positioned in close spatial proximity prior to transcription. In dorsal *svb*-expressing nuclei, the distances between the enhancer probes and the *svb* promoter were further reduced compared with non-expressing nuclei. In ventral *svb*-expressing nuclei, only the *E6*-promoter distance was significantly reduced, whereas the *DG2*-promoter distance showed a modest, non-significant decrease. The distance between the two enhancer probes, *DG2* and *E6*, was also reduced in dorsal *svb*-expressing nuclei but did not reach statistical significance. Because the ventral enhancers *DG3* and *E3* are located adjacent to *DG2* and *E6*, respectively, along the chromosome (Fig. 3a), the *DG2* and *E6* probe signals may also reflect spatial interactions involving these neighboring ventral enhancers.

To further assess whether these pairwise changes reflect broader shifts in three-locus spatial organization, we applied a recently developed probabilistic framework for analyzing DNA-FISH distance distributions^58^. This analysis revealed a significant shift toward shorter inter-probe distances in dorsal *svb*-expressing nuclei (Fig. S2a), suggesting that *svb* expression is associated with increased compaction of the three probed regions within the hub.

To capture the spatial relationship between all three loci, we calculated the centroid distance score of the three DNA-FISH signals, defined as the sum of three-dimensional Euclidean distances of *DG2*, *E6*, and the *svb* promoter from their collective centroid (Fig. 3e). This measurement provides a single readout of how spread out the three loci are around their shared center, with lower values indicating a more compact spatial organization within the nucleus. A threshold of 0.5 µm was applied to define compact clusters. This value exceeds the theoretical minimum centroid distance score of ∼0.43 µm at which all three inter-probe distances simultaneously reach the confocal resolution limit (∼250 nm; centroid distance score = d√3, where d = 0.25 µm), and therefore provides a conservative upper boundary for compact hub conformations. Across all three cell populations, a substantial fraction of nuclei exhibited centroid distance score values below this threshold, with 48% of non-expressing, 56% of dorsal, and 54% of ventral nuclei falling within this compact range (Fig. 3f). In dorsal *svb*-expressing nuclei, the centroid distance score was significantly reduced, resulting in a higher fraction of compact clusters. Ventral *svb*-expressing nuclei also showed a reduction in centroid distance score and a higher fraction of compact clusters, although these differences did not reach statistical significance. No significant difference was observed between dorsal and ventral *svb*-expressing nuclei, indicating that hub compaction is comparable between the two *svb*-expressing populations.

Together, these results show that *svb* regulatory elements frequently occupy compact enhancer-promoter conformations in single nuclei, consistent with the formation of enhancer-promoter hubs. Such hubs are present in both expressing and non-expressing cells, with the highest frequency observed in *svb*-expressing nuclei, indicating that *svb* expression is associated with increased hub compaction.

### The *shavenbaby* enhancer-promoter hub precedes *shavenbaby* expression

The presence of enhancer-promoter hubs in non-expressing nuclei raised the question of whether this organization is established prior to *svb* expression during embryogenesis. To address this, we first examined the chromatin architecture of the *svb* locus across developmental stages.

Analysis of publicly available Micro-C datasets spanning early *D. melanogaster* embryogenesis^59^ revealed that the *svb* regulatory region resides within a well-defined chromatin interaction domain corresponding to the *svb* TAD (Fig. S3). This domain, encompassing the *svb* gene and its associated enhancers, is already apparent in early embryos at nuclear cycles 1-8 (nc1-8), prior to zygotic genome activation, and persists through nc14 and later developmental stages (Fig. S3). While the overall domain structure remains stable, interaction intensity within the domain increases over time, with more pronounced internal contacts emerging at nc14 and persisting at later stages, as reported for other loci^41^. These results indicate that the *svb* regulatory region is embedded within a chromatin interaction domain that is established prior to zygotic genome activation, and that internal contacts within this domain increase before the onset of *svb* expression.

We next asked whether promoter-enhancer interactions are already present prior to *svb* expression. Using UMI-4C, we mapped the genomic interactions of the *svb* promoter in nc14 embryos, a stage preceding *svb* expression (Fig. 4a and Fig. S4). At this stage, the *svb* promoter interacted with multiple distal regions, including known *svb* enhancers, within the *svb* TAD. These interactions were distributed across the locus and resembled those observed in ventral epidermal cells at later stages (Fig. 2e), indicating that promoter-enhancer contacts are already established at nc14.

**Fig. 4.**
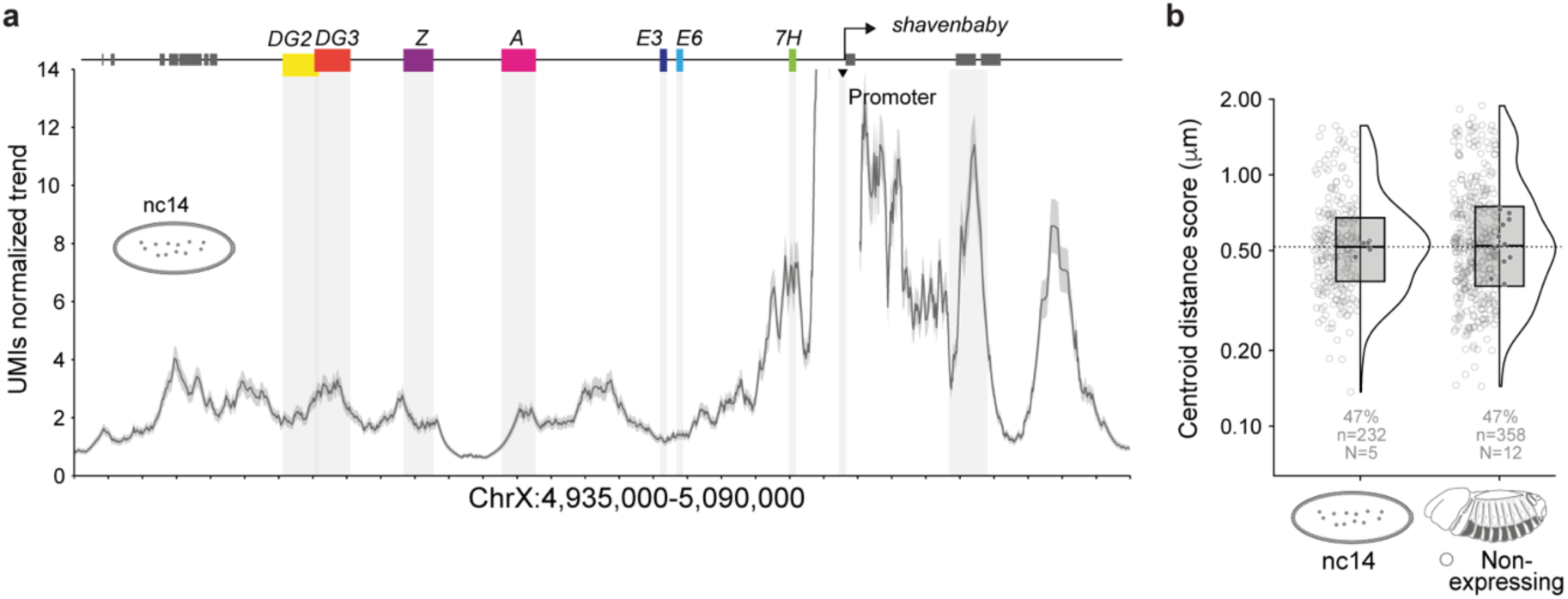
The *shavenbaby* enhancer-promoter hub precedes gene activation (a) UMI-4C interaction profile generated using the endogenous *svb* promoter as the viewpoint in nuclei isolated from nuclear cycle 14 (nc14) embryos. Interaction frequencies are shown as UMI-normalized trends across the *svb* locus. Gray shading indicates the positions of the characterized *svb* enhancers, and the triangle marks the promoter viewpoint. The line represents the mean interaction profile, and the shaded envelope denotes the variability across biological replicates. (b) Centroid distance scores in nc14 nuclei and stage 14 *svb* non-expressing epidermal nuclei. Violin plots, box plots, individual nuclei, embryo-level medians, and statistical testing are presented as in Fig. 3d. The dotted line marks the median centroid distance score of the stage 14 *svb* non-expressing population. Percentages below indicate the fraction of nuclei with centroid distance scores below 0.5 μm, while numbers indicate the number of nuclei analyzed (*n*) and embryos (*N*).

To determine whether these interactions correspond to spatial proximity within individual nuclei, we applied DNA-FISH in nc14 male embryos and calculated the centroid distance score to quantify the spatial relationship between *DG2*, *E6*, and the *svb* promoter (Fig. 4b). Nc14 nuclei exhibited a broad range of conformations, with 47% of nuclei exhibiting centroid distance score values below 0.5 µm (Fig. 4b). The fraction of nuclei with compact conformations was similar in nc14 embryos and stage 14 non-expressing nuclei, indicating that compact enhancer-promoter hubs are already common prior to *svb* expression. Accordingly, analysis of pairwise distance distributions revealed no significant difference between stage 14 non-expressing nuclei and nc14 nuclei (Fig. S2b).

Importantly, the compact conformations observed in early development are unlikely to reflect chromatin condensation, as the *svb* locus lacks repressive histone marks and exhibits open chromatin profiles at this developmental stage (Fig. S5). Thus, the early proximity between the *svb* promoter and distal enhancers occurs within an accessible chromatin environment rather than reflecting broad chromatin repression.

These results suggest that enhancer-promoter hubs at the *svb* locus do not arise de novo at the onset of transcription but instead are already present early in development and become more frequent in expressing nuclei.

### Multiple factors contribute to enhancer-promoter hub organization

So far, our results show that the *svb* regulatory region is organized as an enhancer-promoter hub across multiple cellular contexts. These hubs are present prior to *svb* expression and in non-expressing epidermal cells, and include enhancers regardless of their activity state. We therefore next asked what holds these hubs together. We hypothesized that trans-acting factors, including chromatin architectural proteins and transcription factors, contribute to hub organization.

To identify trans-acting regulators of enhancer-promoter hub organization at the *svb* locus, we first compiled a set of candidate factors based on their expression and binding profiles. We surveyed publicly available *in situ* hybridization datasets^60^ and our cell type-specific RNA-seq data^51^ to identify transcription factors and boundary-associated factors expressed in the embryonic epidermis at stages corresponding to *svb* expression. We then integrated published ChIP-seq datasets^61,62^ to select factors with reproducible binding across the *svb* regulatory region, encompassing its known enhancers (Fig. S6). This approach yielded 12 candidate factors, comprising transcription factors and boundary-associated proteins, all with spatially appropriate expression and reproducible binding across the *svb* regulatory region. In addition, we included three previously characterized regulators of *svb* expression: Arrowhead (Awh)^51^, Pannier (Pnr)^51^ and Ultrabithorax (Ubx)^50^.

To determine whether these factors contribute to the enhancer-promoter hub organization, we downregulated individual candidates using RNAi in the embryonic epidermis and quantified changes in hub structure by multicolor DNA-FISH, as described above (Fig. 5). Hub organization was assessed using the centroid distance score of the *DG2*, *E6*, and promoter DNA-FISH signals, where an increase in centroid distance score reflects reduced proximity between the three loci.

**Fig. 5.**
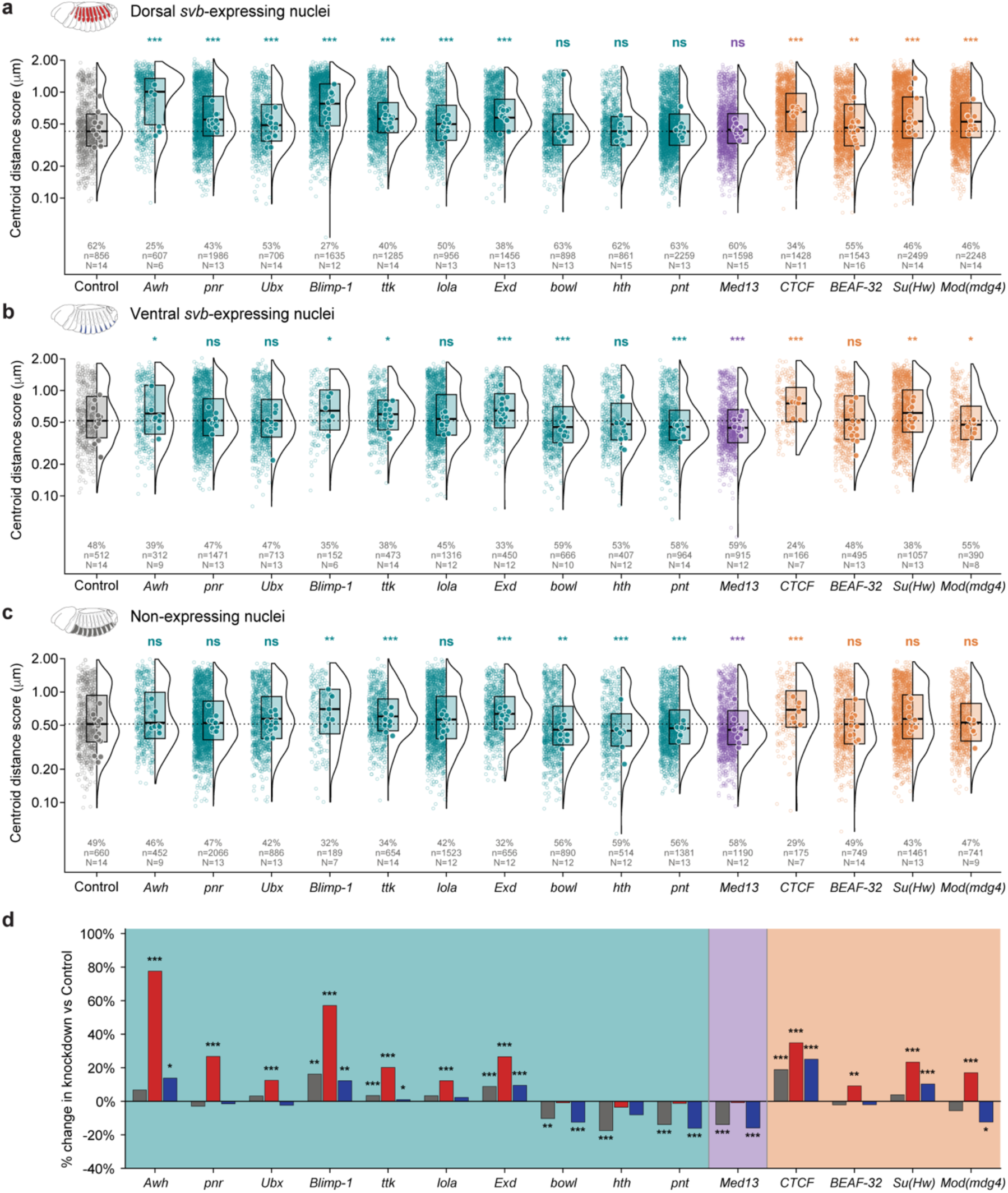
Multiple factors contribute to enhancer-promoter hub organization. (a–c) Centroid distance scores following RNAi-mediated depletion of the indicated transcription factors (teal), Mediator sub-unit (purple) and boundary element proteins (orange) in dorsal *svb*-expressing nuclei (a), ventral *svb*-expressing nuclei (b), and non-expressing nuclei (c). Violin plots, box plots, individual nuclei, and embryo-level medians are presented as in Fig. 3d. The dotted horizontal line indicates the median centroid distance score of the corresponding *LacZ* control (gray). Percentages indicate the fraction of nuclei with centroid distance scores below 0.5 μm, and numbers below indicate the number of nuclei analysed (*n*) and biological replicates (*N*). Statistical significance was assessed relative to *LacZ* controls using two-sided Wilcoxon rank-sum tests with Benjamini-Hochberg correction. ns, not significant; * - *P* < 0.05, ** - *P* < 0.01, *** - *P* < 0.001. (d) Percentage change in mean centroid distance score relative to the corresponding *LacZ* control for each RNAi condition in *svb* non-expressing (gray), dorsal *svb*-expressing (red), and ventral *svb*-expressing nuclei (blue). Positive values indicate reduced hub compaction, whereas negative values indicate increased hub compaction. Asterisks indicate significant differences relative to the corresponding *LacZ* control following Benjamini-Hochberg correction as in (a-c). Background shading distinguishes transcription factor (teal), Mediator sub-unit (purple) and boundary element protein perturbations (orange).

RNAi-mediated downregulation of candidate factors produced factor-and context-specific changes in centroid distance score distributions, reflecting shifts in the fraction of nuclei exhibiting compact enhancer-promoter hubs rather than a complete loss of such conformations. Several perturbations resulted in increased centroid distance score, indicating reduced hub compaction. In dorsal *svb*-expressing nuclei, downregulation of the *svb* regulators *Awh, pnr* and *Ubx*, as well as *Blimp-1*, *ttk*, *lola*, and *Exd*, led to increased centroid distance scores relative to controls, with *Awh* and *Blimp-1* producing the strongest effects (Fig 5a). Similar effects were observed in ventral *svb*-expressing nuclei for a subset of these factors, including *Awh*, *Blimp-1, ttk*, and *Exd* (Fig. 5b). However, the effects of *Awh* and *Blimp-1* downregulation were smaller than those observed in dorsal *svb*-expressing nuclei. In addition, downregulation of *Blimp-1, ttk*, and *Exd* also affected non-expressing epidermal cells (Fig. 5c), indicating that their contribution to hub organization is not restricted to *svb*-expressing nuclei, although the strongest and broadest effects were observed in expressing cell populations.

In contrast, downregulation of a subset of factors resulted in decreased centroid distance score, indicating increased chromatin compaction. These effects were observed for factors such as *bowl*, *hth*, and *Med13*, primarily in ventral and non-expressing nuclei (Fig. 5b-c).

To determine whether chromatin architectural proteins contribute to enhancer-promoter hub organization, we examined boundary-associated factors, including CTCF, BEAF-32, Su(Hw), and their cofactor Mod(mdg4), which are enriched at TAD boundaries and also bind within the *svb* locus (Fig. S6). Downregulation of boundary-associated factors resulted in context-specific changes in centroid distance score distributions, similar to those observed for transcription factors. Depletion of *CTCF* led to increased centroid distance scores in all cell subpopulations, including dorsal and ventral *svb*-expressing and non-expressing nuclei (Fig. 5a-c). In contrast, depletion of *Su(Hw)* and *Mod(mdg4)* affected only *svb*-expressing nuclei, with varying effect sizes, whereas *BEAF-32* depletion led to an increased centroid distance score only in dorsal *svb*-expressing nuclei (Fig. 5a-c).

To test whether these effects reflect alterations in higher-order chromatin organization, we analysed publicly available Hi-C datasets generated following depletion of boundary-associated factors^63^. Despite changes in interaction frequencies, the overall TAD structure at the *svb* locus remained intact (Fig. S7), indicating that depletion of these factors does not disrupt domain organization but rather modulates local chromatin interactions within the locus.

Together, these results indicate that multiple trans-acting factors contribute to enhancer-promoter hub organization at the *svb* locus. The partial and context-specific effects observed upon perturbation of individual factors suggest that hub organization is regulated through combinatorial and context-dependent inputs, and can be modulated independently of higher-order domain structure.

### Perturbations affect hub organization either broadly or through specific pairwise relationships

The centroid distance score identified perturbations that altered overall compaction of the *svb* enhancer-promoter hub but did not reveal whether these changes reflected hub-wide shifts or changes in specific pairwise relationships. To address this, we analyzed the pairwise distance distributions between *DG2*, *E6*, and the *svb* promoter using the same probabilistic framework^58^ described above, comparing each RNAi condition to its corresponding control population (Fig. S8).

This analysis showed that most perturbations that increased centroid distance score were accompanied by changes across multiple pairwise distances, consistent with broad reorganization of the enhancer-promoter hub. Among transcription factors, RNAi-mediated downregulation of *Exd* altered all three pairwise relationships across dorsal, ventral, and non-expressing nuclei (Fig. S8a). Similarly, downregulation of *Awh*, *pnr*, *Ubx*, *Blimp-1*, *ttk*, and *lola* altered all three pairwise relationships in dorsal *svb*-expressing nuclei but produced more selective effects in ventral and non-expressing nuclei (Fig. S8a).

Boundary-associated factors showed a similar range of effects. RNAi-mediated downregulation of *CTCF* altered all three pairwise relationships in every cell population examined, whereas downregulation of *Su(Hw)*, *BEAF-32*, and *Mod(mdg4)* produced more context-specific changes that were restricted to particular cell populations or pairwise relationships (Fig. S8b).

Together, these analyses indicate that many trans-acting factors affect the *svb* hub through hub-wide shifts in the spatial organization of the three regions, while others influence more specific spatial relationships within the hub. Thus, individual perturbations can alter hub organization at multiple levels, rather than simply disrupting a single enhancer-promoter contact.

### Perturbation of enhancer-promoter hub organization is associated with reduced *shavenbaby*-dependent phenotypes

The analyses above showed that downregulation of different trans-regulators alters the organization of the *svb* enhancer-promoter hub in a cell type-specific manner. We next asked how these changes in hub organization affect *svb*-dependent phenotypic outputs. To address this, we quantified dorsal and ventral trichome numbers in first instar larvae following RNAi-mediated downregulation of factors that affect hub organization (Fig. 6 and Fig. S9). Trichome number serves as a direct readout of *svb* function in the epidermis^2^.

**Fig. 6.**
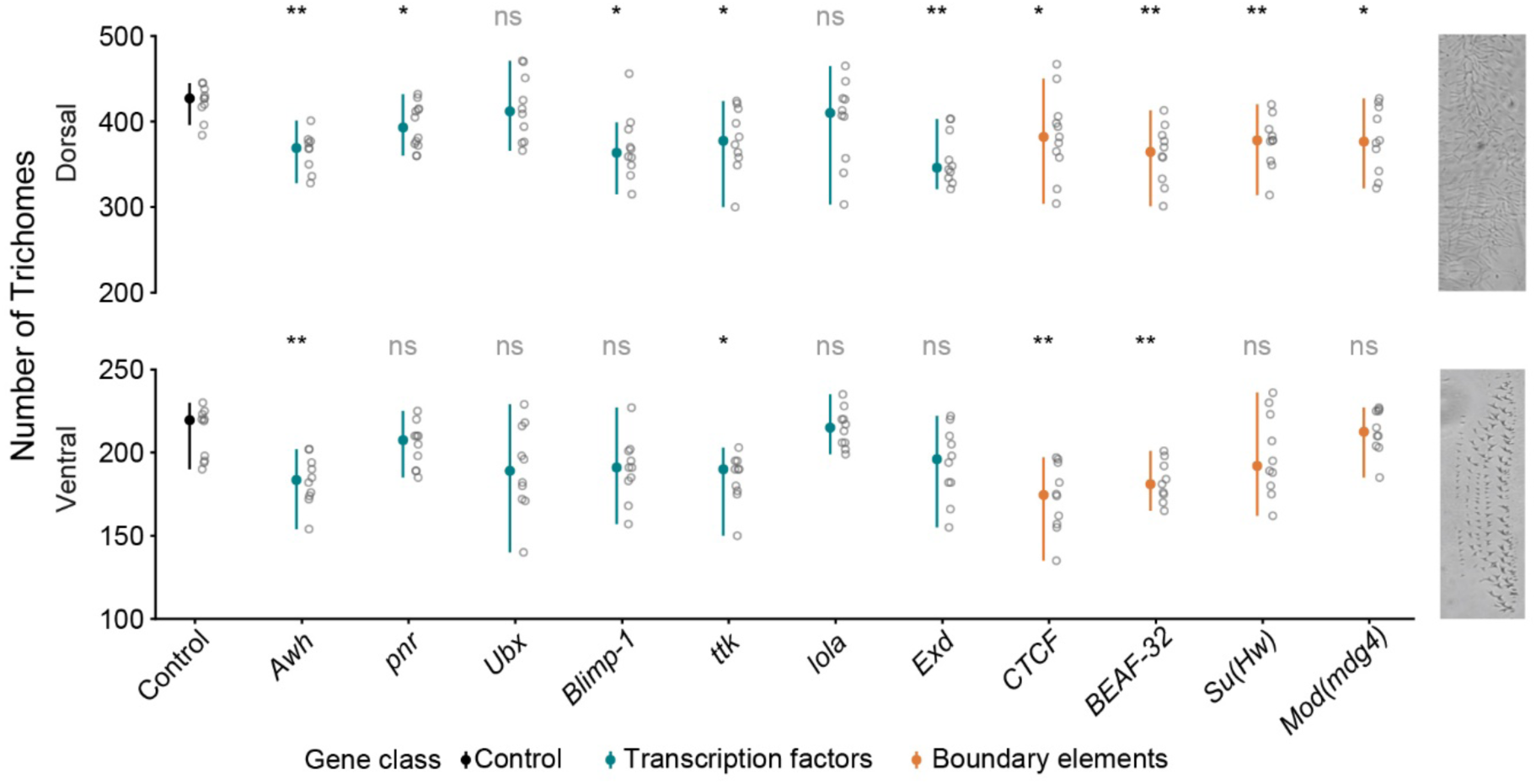
Perturbation of factors contributing to enhancer-promoter hub organization affects *svb*-dependent trichome formation Number of trichomes in the dorsal (top) and ventral (bottom) cuticle of the fifth abdominal segment of first instar larvae following RNAi-mediated depletion of the indicated transcription factors (teal) and boundary element proteins (orange) compared with the *LacZ* control (black). Open circles indicate individual larvae, colored circles indicate the mean, and vertical lines denote standard deviation. Statistical significance was assessed relative to the *LacZ* control using two-sided statistical tests with Benjamini–Hochberg correction. ns, not significant; * - *P* < 0.05, ** - *P* < 0.01, *** - *P* < 0.001. Representative cuticle images are shown on the right.

As expected, downregulation of the *svb* regulators *Awh* and *pnr* significantly reduced the number of dorsal trichomes, consistent with their function as activators of the *svb E6* enhancer (Fig. 6). Interestingly, downregulation of *Awh* also reduced the number of ventral trichomes, consistent with its effect on hub organization in ventral epidermal cells. In addition to these factors, downregulation of several other regulators reduced trichome number in the same domains in which they affected hub compaction, although the magnitude and spatial distribution of these effects varied between factors. For example, knockdown of *ttk*, *Exd*, and *CTCF* reduced trichome numbers in both dorsal and ventral domains, consistent with their effects on hub compaction in these regions (Fig. 6).

Downregulation of *Blimp-1* and the boundary-associated factors *su(Hw)*, *BEAF-32*, and *Mod(mdg4)* primarily affected dorsal trichomes, consistent with their stronger effects on hub organization in dorsal epidermis (Fig. 6). In these cases, the spatial distribution of the phenotypic effects closely matched the domains in which these factors influenced hub organization, indicating that loss of a compact enhancer-promoter hub is frequently associated with reduced *svb*-dependent output.

Conversely, knockdown of *lola* and *Ubx* altered hub organization relative to control without producing significant changes in dorsal trichome number, although *Ubx* showed modest effects in ventral regions (Fig. 6). These observations indicate that changes in enhancer-promoter hub organization are not always associated with altered *svb* functions.

Together, these findings establish a functional link between *svb* enhancer-promoter hub organization and trichome formation. Disruption of hub organization was frequently associated with reduced *svb*-dependent trichome formation in the corresponding epidermal domains. However, individual perturbations produced only partial effects on both hub organization and trichome formation, suggesting that multiple factors contribute to hub formation and *svb* function across the embryonic epidermis.

## Discussion

In this study, we showed that the *svb* regulatory region is organized as a multi-enhancer-promoter hub. This hub is present before *svb* expression and in epidermal cells that do not express *svb* and includes enhancers regardless of their activity state. However, compact enhancer-promoter hubs are more frequent in *svb*-expressing cells, consistent with a role in transcriptional activation. We further show that hub organization does not depend on a single factor. Instead, the contribution of individual trans-regulators to the 3D organization of the locus varies between epidermal contexts, suggesting that similar enhancer-promoter hubs can be maintained through different combinations of regulatory inputs. Finally, changes in hub organization are frequently associated with changes in *svb*-dependent trichome patterning, providing evidence for a functional link between enhancer-promoter hub organization and the developmental outcome of *svb* expression. Together, these findings suggest that enhancer-promoter hubs enable diverse trans-regulatory environments to generate precise developmental gene expression through a shared three-dimensional organization.

Enhancer-promoter hubs have been documented at several other developmental loci. One of the early examples of such organization was described at the mouse *HoxD* gene cluster, where the 5′ *Hoxd* genes are regulated by multiple distal enhancers distributed across a large gene desert that together form a “regulatory archipelago”^25^. In developing digits, the active gene cluster contacts several of these regulatory elements, whereas in non-expressing tissue a subset of these contacts is already present, forming a less elaborate ground-state organization^25^. Multi-way analyses at the α-globin locus also showed that multiple regulatory elements and promoters can participate simultaneously in a shared chromatin hub rather than interacting exclusively^64^. High-throughput 3C-based studies have further identified multiple loci organized as multi-enhancer-promoter hubs in developing embryos^41,42^, supporting the idea that such organization is a recurring feature of developmental gene regulation. However, while these studies have largely defined the architecture of such hubs, the mechanisms that shape their organization remain poorly understood.

What factors contribute to the establishment and maintenance of enhancer-promoter hubs? Our perturbation experiments indicate that enhancer-promoter hub organization depends on the combined contributions of multiple chromatin architectural proteins and transcriptional regulators with different effects across developmental contexts. One possible model is that chromatin architectural proteins help establish or stabilize enhancer-promoter proximity within a landscape of accessible chromatin early in embryogenesis, independently of context-specific transcription factors. This model is supported by a recent study showing that enhancer-promoter interactions that form stably across multiple tissues during mouse embryogenesis are frequently associated with CTCF binding^42^. Consistent with this idea, CTCF contributed to hub organization across all epidermal contexts examined, including in non-expressing cells, whereas other architectural proteins showed more context-specific effects.

As development proceeds, trans-regulatory environments become increasingly complex, even within a single tissue such as the epidermis. Transcription factors expressed in different epidermal domains may bind the *svb* regulatory region and increase the frequency or stability of compact hubs. Consistent with this idea, downregulation of multiple transcription factors affected hub organization in a context-dependent manner. These included both known regulators of *svb* and factors not previously implicated in *svb* regulation. Among the latter, *Blimp-1* was notable because its depletion caused the strongest effect on hub organization in dorsal epidermal cells. Interestingly, Blimp-1 motifs are enriched near promoters engaged in enhancer-promoter interactions in differentiating neurons^41^, suggesting that Blimp-1 may also contribute to enhancer-promoter organization in other developmental contexts. These findings raise the possibility that enhancer-promoter hubs with similar spatial organization are maintained by partially different combinations of regulatory proteins and therefore differ in their molecular composition across epidermal contexts.

The spatial proximity of enhancers and the promoter may facilitate transcriptional activation once the appropriate transcription factors bind their target enhancers. These hubs may also act as structural scaffolds that facilitate the assembly of transcriptional hubs enriched in trans-regulators and transcriptional machinery. The relationship between enhancer-promoter hubs and transcriptional hubs may be reciprocal, as the assembly of transcriptional machinery could further stabilize enhancer-promoter proximity or reinforce interactions within the hub. Consistent with this model, live imaging of the *Sox2* locus showed that the *Sox2* control region and promoter tend to stably co-localize, while transcriptional burst size and frequency increase upon dynamic encounters between RNA polymerase II or Mediator condensates and these pre-positioned regulatory elements^30^. Because our DNA-FISH measurements capture fixed snapshots, they cannot determine whether compact hubs precede transcriptional hub assembly, are stabilized by it, or both. Future live imaging of the *svb* locus in developing embryos will determine the relationship between enhancer-promoter hubs, transcriptional hubs, and active transcription.

The organization of the *svb* locus may also contribute to the robustness of its developmental output^2^. Previous work showed that a BAC containing the complete *svb cis*-regulatory region, inserted on another chromosome, co-localizes with the endogenous *svb* locus and can compensate for deletions of endogenous enhancers that otherwise reduce phenotypic robustness at elevated temperature^52^. This co-localization restored *svb* transcriptional output and trichome formation under stressful conditions, leading the authors to propose that multiple *svb* enhancers contribute to shared transcriptional hubs that buffer against environmental perturbations. They further proposed that enhancer-enhancer interactions may provide a structural basis for these hubs, allowing multiple enhancers to reinforce a shared transcriptional environment^52^. Our findings support this prediction and provides direct evidence that the endogenous regulatory region is organized as a multi-enhancer-promoter hub. Moreover, this hub persists across epidermal domains despite differences in enhancer activity and in the trans-regulators that contribute to its organization. Thus, the robustness of the *svb* system may arise from both the spatial integration of multiple enhancers and the ability of different combinations of regulatory proteins to maintain the hub across developmental contexts. Notably, not all perturbations that altered hub organization produced detectable changes in trichome patterning. For example, downregulation of *lola* and *Ubx* altered hub organization without significantly affecting trichome number, suggesting that modest changes in hub organization may not necessarily translate into measurable developmental phenotypes.

Together, our findings suggest that developmental genes regulated by multiple enhancers rely on a shared three-dimensional regulatory organization that is maintained across different developmental contexts. More broadly, multi-enhancer-promoter hubs may represent a general principle of developmental gene regulation, integrating multiple regulatory inputs within a common spatial assembly to balance robustness with regulatory flexibility.

## Limitations

Several limitations should be considered when interpreting these findings. UMI-4C reflects population-averaged pairwise interaction frequencies and cannot resolve whether enhancer-promoter contacts co-occur simultaneously within individual nuclei. DNA-FISH provides single-cell resolution but visualizes only three genomic loci simultaneously, therefore, our measurements of the promoter and two enhancers likely represent a subset of the full multi-enhancer hub. Multiplexed imaging or multi-contact chromatin conformation approaches will be required to resolve the complete composition of the hub.

RNAi-mediated knockdown is inherently incomplete, and penetrance can vary between RNAi lines. We selected RNAi lines that have been functionally validated in previous studies. Additionally, because RNAi was induced in post-gastrulation epidermal cells, the contribution of these factors to hub establishment at earlier developmental stages remains unaddressed. Although all candidates bind the *svb* regulatory region, some observed effects on hub organization may nevertheless be indirect.

Finally, because all analyses were performed in fixed embryos, they provide only static snapshots of enhancer-promoter organization. Future live-cell imaging approaches will be required to determine how hub organization changes over time and how these changes relate to transcriptional activity during development.

## Methods

### Transgenic Constructs and Fly Strains

Enhancer fragments were amplified by PCR from genomic DNA of wild-type *D. melanogaster* (*Oregon^R^*) and cloned into the reporter constructs *placZattB, pS3AR* and *pS3AG* using Gibson Assembly (see Table S1 for details). Reporter constructs were integrated into separated *attP* landing sites by Rainbow Transgenic Flies and then combined to generate the two tester lines: *DG2B::dsRed; Z1.3L::lacZ; E6B::GFP* and *E3::dsRed; 7H::lacZ*.

Additional strains used were: *Oregon^R^, P{w[+mC]=Sxl-Pe-EGFP.G}G5b, w[*]* and *w;P{y[+t7.7] w[+mC]=GMR43B11-GAL4}attP2*^65^. All lines used in this study, including RNAi lines, are summarized in Supplementary Table S2.

### Embryo Staining and imaging

Stage 15 embryos were collected, fixed and stained using standard protocols with mouse anti-βGal (1:500, Promega), rabbit anti-RFP (1:500, MBL), chicken anti-GFP (1:500, Ave Laboratories), AlexaFluor 488 goat anti-mouse, AlexaFluor 488 goat anti-chicken, AlexaFluor 555 goat anti-rabbit, and AlexaFluor 633 goat anti-mouse (1:500, Invitrogen) antibodies. Stained embryos were imaged on a ZEISS LSM 900 Confocal Microscope. Images were processed using Fiji (https://imagej.net/)^66^ as previously described^56^.

### Cell type-specific UMI-4C

Embryos from tester lines (*DG2B::dsRed; Z1.3L::lacZ; E6B::GFP* or *E3::dsRed; 7H::lacZ*) or wild-type (*Oregon^R^*) embryos were collected on grape-juice plates in large population cages (Flystaff Cat # 59-104) for 2 h and aged to the appropriate stage at 25°C (Table S3). Aged embryos were crosslinked in 1.8% formaldehyde for 20 min, quenched, flash-frozen in liquid nitrogen, and stored at −80°C until processing. Nuclei from ∼500 mg flash-frozen embryos were isolated and immunostained with appropriate antibodies (Table S3) according to the BiTS-ChIP (Batch Isolation of Tissue-Specific Chromatin for Immunoprecipitation) protocol^57^. Enhancer-specific cell populations were sorted on Becton-Dickinson FACSAria IIIu (gating strategy is shown in Fig. S10). Sorting purity (>95%) was confirmed by re-analysis of an aliquot before proceeding. Sorted nuclei were collected into 1ml of chilled PBTB buffer and processed immediately for UMI-4C library preparation.

UMI-4C libraries were prepared independently from 1 × 10⁶ sorted crosslinked nuclei per population across three to four independent biological replicates (separate embryo collections and FACS sorts). Chromatin was permeabilized with 0.1 % SDS, quenched with Triton X-100, and digested with MseI (100 U, 3 h at 37°C). Proximity ligation was performed overnight at 16°C under dilute conditions (T4 DNA ligase, 2,000 U); crosslinks were reversed, RNA removed by RNase A, and 3C DNA purified by phenol-chloroform extraction and ethanol precipitation. 3C DNA was sonicated to ∼150–700 bp (Covaris), end-repaired, A-tailed, and adapter-ligated (NEBNext Ultra II). After USER® enzyme treatment and ssDNA generation by heat denaturation, libraries were amplified in two nested PCR rounds using viewpoint-specific primers and unique i7 index primers (primer sequences in Table S4).

Libraries were sequenced on DNBSEQ-G400 (PE150) by Syntezza Bioscience Ltd. Each viewpoint was sequenced to a target depth of ∼1 × 10⁶ mapped reads per library; three to four libraries per condition were pooled to increase complexity. Data were processed using the UMI4Cats R package (v1.0.0) ^67^ with Bowtie2 (v2.4.5) alignment to dm6: reads were demultiplexed by viewpoint DS primer, trimmed, aligned, and UMI-collapsed to remove PCR duplicates. Interaction profiles were computed within a 2 Mb window around each viewpoint (*makeUMI4C*; *groupsUMI4C*), smoothed by adaptive Gaussian kernel (*sd* = 1), and visualized with enhancer and gene annotations using ggplot2 (v3.4.0).

### 3D multicolor DNA-FISH

#### Probe synthesis

Fluorescent probes targeting the *svb* promoter (Alexa Fluor 488), *E6* (Atto 550), and *DG2* (DyLight 635) loci (3-5 kb each; Table S4) were amplified from genomic DNA, cloned into pGEM®-T Easy, sequence-verified, and re-amplified with M13 primers. Probes were directly labelled by nick translation using aminoallyl-dUTP-conjugated fluorophores (15°C, 8–16 h; confirmed by 150–500 bp smear on agarose gel) and resuspended in hybridization buffer (HyB - 50% formamide, 5× SSC, 0.3% CHAPS, 100 µg/ml heparin, 0.1 mg/ml salmon sperm DNA).

#### Embryo processing and imaging

DNA-FISH was performed specifically on male embryos, to make sure that each nuclei contain one allele of the X-linked *svb* locus. To enable sex-specific collection of male embryos across all conditions, females of the *Sxl-Pe-eGFP* reporter line were used in all crosses; male embryos were identified by absence of *Sxl-Pe-eGFP* fluorescence (>98% accuracy) and confirmed by midgut autofluorescence. Age-appropriate male embryos were fixed in 1.8% formaldehyde according to standard protocols and stored in 100% methanol at −20°C until use.

The embryos stored in methanol were rehydrated sequentially in freshly prepared methanol/PBT solutions (70%, 50%, and 30%) for 5 min each on a nutator, followed by equilibration in 100% PBT for 1 h. To gradually transition samples into hybridization conditions, PBT was stepwise replaced with increasing concentrations of formamide in 2× SSCT (80% PBT/20% formamide and 50% PBT/50% formamide, 20 min each), followed by incubation in 100% HyB for 1 h on a rotator. Embryonic DNA was denatured in 100 μl HyB at 82°C for 12 min in a thermomixer. In parallel, 10 μl of fluorescently labelled DNA probe diluted in 30 μl HyB was denatured at 90°C for 10 min, cooled to 82°C and added directly to the embryos at 82°C. Hybridization was carried out overnight (14-20 h) at 37°C with gentle agitation (450-500 rpm) in the dark. Following hybridization, embryos were washed twice in 50% formamide in 2× SSCT at 37°C for 1 h, once in 20% formamide in 2× SSCT for 20 min, and three times in 2× SSCT for 10 min each. Embryos were then stained with DAPI (1:1000 dilution of a 1 mg/mL stock) in PBT for 10 min, followed by a final wash in PBT for 10 min. Samples were transferred into 1× PBS (≈50 μl remaining volume) and mounted in 40 μl ProLong™ Gold Antifade Mountant. Slides were allowed to cure overnight at 4°C prior to imaging. Throughout the procedure, embryos were kept hydrated and protected from light to preserve nuclear integrity and fluorophore stability.

Images were acquired on a Zeiss LSM 900 with a Plan-Apochromat 63×/1.40 NA oil-immersion objective and Airyscan 2 detector in 3D super-resolution mode using sequential excitation; all acquisition parameters were held constant within batches (Table S5). Z-stacks of ∼61 sections spanning 8.4 µm were acquired (voxel: 0.0495 × 0.0495 × 0.140 µm) and processed using Zeiss ZEN (v3.5) Airyscan 3D automatic reconstruction. Chromatic shifts between channels were corrected using 0.1 µm TetraSpeck™ microspheres (Thermo Fisher Scientific) imaged under identical optical conditions; a global linear 3D transformation per non-reference channel was calculated by 3D Gaussian bead centroid fitting in the Zeiss ZEN Channel Alignment (Extended) module (633 nm reference) and applied to all experimental datasets.

### Data analysis

Spots were detected using the RS-FISH plugin (Fiji/ImageJ v2.9.0; Advanced mode; DoG detection, σ = 1.8, anisotropy factor 0.81, RANSAC fitting) applied via a custom batch macro identically across all images. Nuclei were segmented from the DAPI channel using Cellpose (detailed in Supplementary Method S1). Subpixel spot coordinates from the three channels were assembled into 3D triplets in Python (v3.10): for each channel, the top 8,000 spots (ranked by RS-FISH intensity, chosen to exceed the expected number of allelic signals per field while limiting spurious matches in dense nuclear fields) were retained and assigned to a nucleus by direct voxel-indexed lookup into the corresponding segmentation mask, with spots not falling within a segmented nucleus excluded from further analysis. For every nucleus containing at least one detected spot in all three channels (*svb* promoter, *E6*, *DG2*), a triplet was accepted only if each of the three pairwise 3D distances, computed in physical units (µm) using the voxel calibration specific to that acquisition, fell within an empirically determined cutoff of ≤1.2 µm, determined from the inflection point of the pairwise nearest-neighbor distance distribution across all detected spots; this cutoff is well below the nuclear diameter of nc14 epidermal cells (∼8-10 µm) and, combined with the nucleus-restriction step above, ensures that all three signals within a triplet originate from the same nucleus. Nuclei yielding more than two valid triplets were flagged for manual inspection and excluded from the analysis. Physical voxel dimensions were extracted directly from the acquisition metadata of each raw CZI file, all pixel-to-micron distance conversions were performed using the calibration specific to each embryo’s own acquisition. Triplets were restricted to manually defined epidermal 2D ROIs drawn in Fiji/ImageJ by an observer blinded to genotype. Dorsal and ventral epidermal boundaries were delineated based on *svb* mRNA expression detected by HCR-RNA FISH (Fig. 1a), which marks the two spatially distinct *svb*-expressing epidermal populations. Non-expressing (NE) ROIs were drawn in the intervening region confirmed to lack *svb* transcript signal. ROI polygons were imported into Python using the roifile library, converted to Shapely (v2.0) polygons, and triplets retained only if all three loci fell within the ROI in the x–y plane. ROI-filtered tables were mapped to three nuclear populations: *svb* non-expressing (NE), dorsal *svb*-expressing (D), and ventral *svb*-expressing (V) nuclei.

### Centroid distance score and statistical analysis

Hub compactness for each nucleus was quantified by the centroid distance score (µm), defined as the sum of the 3D Euclidean distances from each of the three DNA-FISH loci to their shared geometric centroid. Let *p*₁ = (x₁, y₁, z₁), *p*₂ = (x₂, y₂, z₂), and *p*₃ = (x₃, y₃, z₃) denote the subpixel coordinates (µm) of the *svb* promoter, *E6*, and *DG2* loci respectively, and let their geometric centroid be:

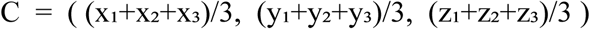

The centroid distance score is then:

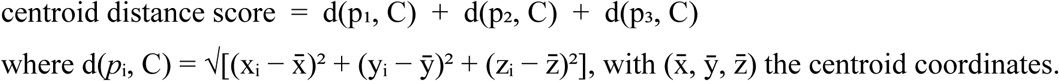

where d(*p*ᵢ, C) = √[(xᵢ − x^-^)² + (yᵢ − ȳ)² + (zᵢ − z^-^)²], with (x^-^, ȳ, z^-^) the centroid coordinates.

A lower centroid distance score indicates greater spatial compaction of the *svb* promoter-*E6*-*DG2* hub. Triplets with non-finite or zero centroid distance score values were excluded.

Statistical analyses were performed in R (v4.3.1). The sample unit *n* is defined as individual nuclei (triplets), with embryo-level medians overlaid to account for between-embryo variation. Centroid distance score distributions in dorsal and ventral *svb*-expressing nuclei were compared independently to non-expressing nuclei per gene using two-sided Wilcoxon rank-sum tests at a significance threshold of α = 0.05; p-values were Benjamini–Hochberg corrected across all comparisons within each gene. Adjusted p-values are reported as: * p < 0.05, ** p < 0.01, *** p < 0.001. Results were visualized in ggplot2 (v3.4.0) with half-violin plots, per-nucleus jitter, and embryo-level median overlays with gene-class background shading.

### Pairwise distance analysis

For each accepted probe triplet, the three pairwise 3D Euclidean distances (*svb* promoter-*E6*, *svb* promoter-*DG2*, and *E6*-*DG2*) were computed. Embryo-condition groups containing fewer than 8 triplets were excluded prior to statistical comparison. Pairwise distance distributions in dorsal and ventral *svb*-expressing nuclei were each compared independently to non-expressing nuclei using two-sided Wilcoxon rank-sum tests at the nucleus/triplet level, with p-values Benjamini-Hochberg corrected across the two comparisons within each probe-pair group (* p ≤ 0.05, ** p ≤ 0.01, *** p ≤ 0.001, **** p ≤ 0.0001; ns, not significant).

Pairwise distance distributions analysis was performed according to Le et al. 2026^58^ (see supplementary method for more information).

### RNAi analysis of candidate *svb* enhancer-promoter hub regulators

To identify regulators of *svb* enhancer-promoter hub organization, we examined the effects of RNAi-mediated downregulation of 15 candidate regulators. UAS-RNAi males (30-40) were crossed to 100-120 virgin females carrying *Sxl-Pe*-*eGFP; Kni-Gal4* at 25°C. Male embryos were collected and processed as described above for DNA-FISH. The effect of each RNAi line on hub organization was determined by comparing the centroid distance score in each epidermal cell subpopulation with that of control embryos obtained from crosses to a *UAS-LacZ* line. The percentage change relative to the control was calculated separately for each epidermal subpopulation as:

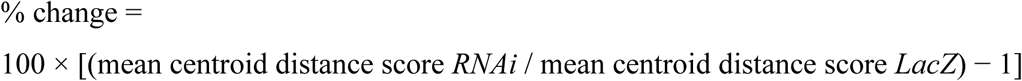

Positive values indicate reduced hub frequency relative to control, whereas negative values indicate increased compaction. Statistical significance for each RNAi line relative to the *LacZ* control was assessed using two-sided Wilcoxon rank-sum tests on per-nucleus centroid distance score values within each subpopulation, with p-values Benjamini–Hochberg corrected across all genes tested within that subpopulation (* p ≤ 0.05, ** p ≤ 0.01, *** p ≤ 0.001).

### Cuticle preparation and trichome quantification

First-instar larvae from RNAi crosses were heat-treated at 65°C for 5 min, mounted in lactic acid, and cleared at 60°C for ≥1 h under light pressure. Brightfield images were acquired on a Zeiss Axio Vert.A1 (Plan-Neofluar 40×/0.75 NA; Axiocam 807 mono). Trichomes were counted on the dorsal and ventral domains of abdominal segment A5 using the Fiji/ImageJ Cell Counter plugin.

## Acknowledgments

We thank Nicolás Frankel and Ramon Birenbaum for critical comments that improved the manuscript. We thank Noam Kaplan, Yoav Shechtman and members of the Preger-Ben Noon lab for helpful discussions. We thank OpenAI’s ChatGPT (GPT-5.5, 2026) for assistance with language editing of the manuscript. This work was supported by a grant from the Israel Science Foundation (No. 2567/20) and by a Short-Term Personal Grant from The Rappaport Family Institute for Research in the Bio Medical Sciences to E.P.B.N. and Gelman-Lazar Research Grant awarded to S.Y.N.

## Author contributions

Conceptualization: E.P.B.N., S.Y.N.

Methodology: S.Y.N.

Investigation: S.Y.N., S.R.

Supervision: E.P.B.N.

Writing: E.P.B.N., S.Y.N.

## Material & Correspondence

All material are available upon request from Ella Preger-Ben Noon.

## References

1. Levine, M. Transcriptional enhancers in animal development and evolution. Curr. Biol. 20, R754–63 (2010).

2. Frankel, N., Davis, G. K., Vargas, D., Wang, S., Payre, F. & Stern, D. L. Phenotypic robustness conferred by apparently redundant transcriptional enhancers. Nature 466, 490–3 (2010).

3. Perry, M. W., Boettiger, A. N., Bothma, J. P. & Levine, M. Shadow Enhancers Foster Robustness of Drosophila Gastrulation. Current Biology vol. 20 (2010).

4. Osterwalder, M., Barozzi, I., Tissiéres, V., Fukuda-Yuzawa, Y., Mannion, B. J., Afzal, S. Y., Lee, E. A., Zhu, Y., Plajzer-Frick, I., Pickle, C. S., Kato, M., Garvin, T. H., Pham, Q. T., Harrington, A. N., Akiyama, J. A., Afzal, V., Lopez-Rios, J., Dickel, D. E., Visel, A. & Pennacchio, L. A. Enhancer redundancy provides phenotypic robustness in mammalian development. Nature 554, (2018).

5. Kvon, E. Z., Waymack, R., Gad, M. & Wunderlich, Z. Enhancer redundancy in development and disease. Nat. Rev. Genet. 22, 324–336 (2021).

6. Schoenfelder, S. & Fraser, P. Long-range enhancer–promoter contacts in gene expression control. Nat. Rev. Genet. 1 (2019) doi:10.1038/s41576-019-0128-0.

7. Furlong, E. E. M. & Levine, M. Developmental enhancers and chromosome topology. Science (1979). 361, 1341–1345 (2018).

8. Levine, M., Cattoglio, C. & Tjian, R. Looping back to leap forward: transcription enters a new era. Cell 157, 13–25 (2014).

9. Sanyal, A., Lajoie, B. R., Jain, G. & Dekker, J. The long-range interaction landscape of gene promoters. Nature 489, 109–113 (2012).

10. Goel, V. Y., Huseyin, M. K. & Hansen, A. S. Region Capture Micro-C reveals coalescence of enhancers and promoters into nested microcompartments. Nat. Genet. 55, 1048–1056 (2023).

11. Zhang, Y., Wong, C.-H., Birnbaum, R. Y., Li, G., Favaro, R., Ngan, C. Y., Lim, J., Tai, E., Poh, H. M., Wong, E., Mulawadi, F. H., Sung, W.-K., Nicolis, S., Ahituv, N., Ruan, Y. & Wei, C.-L. Chromatin connectivity maps reveal dynamic promoter–enhancer long-range associations. Nature 504, 306–310 (2013).

12. Ghavi-Helm, Y., Klein, F. A., Pakozdi, T., Ciglar, L., Noordermeer, D., Huber, W. & Furlong, E. E. M. Enhancer loops appear stable during development and are associated with paused polymerase. Nature 512, 96–100 (2014).

13. Jin, F., Li, Y., Dixon, J. R., Selvaraj, S., Ye, Z., Lee, A. Y., Yen, C.-A., Schmitt, A. D., Espinoza, C. A. & Ren, B. A high-resolution map of the three-dimensional chromatin interactome in human cells. Nature 503, 290–294 (2013).

14. Kragesteen, B. K., Spielmann, M., Paliou, C., Heinrich, V., Schöpflin, R., Esposito, A., Annunziatella, C., Bianco, S., Chiariello, A. M., Jerković, I., Harabula, I., Guckelberger, P., Pechstein, M., Wittler, L., Chan, W.-L., Franke, M., Lupiáñez, D. G., Kraft, K., Timmermann, B., Vingron, M., Visel, A., Nicodemi, M., Mundlos, S. & Andrey, G. Dynamic 3D chromatin architecture contributes to enhancer specificity and limb morphogenesis. Nat. Genet. 1 (2018) doi:10.1038/s41588-018-0221-x.

15. Gschwind, A. R., Mualim, K. S., Karbalayghareh, A., Sheth, M. U., Dey, K. K., Jagoda, E., Nurtdinov, R. N., Xi, W., Tan, A. S., Galante, J., Jones, H., Ma, X. R., Yao, D., Amgalan, D., Ray, J., Munger, C. J., Nasser, J., Avsec, Ž., James, B. T., Shamim, M. S., Durand, N. C., Rao, S. S. P., Mahajan, R., Doughty, B. R., Andreeva, K., Ulirsch, J. C., Fan, K., Perez, E. M., Nguyen, T. C., Kelley, D. R., Finucane, H. K., Moore, J. E., Weng, Z., Kellis, M., Bassik, M. C., Ustun, B., Price, A. L., Beer, M. A., Guigó, R., Stamatoyannopoulos, J. A., Lieberman Aiden, E., Greenleaf, W. J., Leslie, C. S., Steinmetz, L. M., Kundaje, A. & Engreitz, J. M. An encyclopedia of human enhancer–gene regulatory interactions. Nature (2026) doi:10.1038/s41586-026-10781-4.

16. Uyehara, C. M. & Apostolou, E. 3D enhancer-promoter interactions and multi-connected hubs: Organizational principles and functional roles. Cell Rep. 42, 112068 (2023).

17. Perlman, B. S., Burget, N., Zhou, Y., Schwartz, G. W., Petrovic, J., Modrusan, Z. & Faryabi, R. B. Enhancer-promoter hubs organize transcriptional networks promoting oncogenesis and drug resistance. Nat. Commun. 15, 8070 (2024).

18. Zhao, J. & Faryabi, R. B. Spatial promoter–enhancer hubs in cancer: organization, regulation, and function. Trends Cancer 9, 1069–1084 (2023).

19. Zhao, J., Zhou, Y., Tzelepis, I., Burget, N. G., Shi, J. & Faryabi, R. B. Oncogenic transcription factors instruct promoter-enhancer hubs in individual triple negative breast cancer cells. Sci. Adv. 10, (2024).

20. Beagrie, R. A., Scialdone, A., Schueler, M., Kraemer, D. C. A., Chotalia, M., Xie, S. Q., Barbieri, M., de Santiago, I., Lavitas, L.-M., Branco, M. R., Fraser, J., Dostie, J., Game, L., Dillon, N., Edwards, P. A. W., Nicodemi, M. & Pombo, A. Complex multi-enhancer contacts captured by genome architecture mapping. Nature 543, 519–524 (2017).

21. Oudelaar, A. M., Davies, J. O. J., Hanssen, L. L. P., Telenius, J. M., Schwessinger, R., Liu, Y., Brown, J. M., Downes, D. J., Chiariello, A. M., Bianco, S., Nicodemi, M., Buckle, V. J., Dekker, J., Higgs, D. R. & Hughes, J. R. Single-allele chromatin interactions identify regulatory hubs in dynamic compartmentalized domains. Nat. Genet. 50, 1744–1751 (2018).

22. Chandra, A., Yoon, S., Michieletto, M. F., Goldman, N., Ferrari, E. K., Abedi, M., Johnson, I., Fasolino, M., Pham, K., Joannas, L., Kee, B. L., Henao-Mejia, J. & Vahedi, G. Quantitative control of Ets1 dosage by a multi-enhancer hub promotes Th1 cell differentiation and protects from allergic inflammation. Immunity 56, 1451–1467.e12 (2023).

23. Allahyar, A., Vermeulen, C., Bouwman, B. A. M., Krijger, P. H. L., Verstegen, M. J. A. M., Geeven, G., van Kranenburg, M., Pieterse, M., Straver, R., Haarhuis, J. H. I., Jalink, K., Teunissen, H., Renkens, I. J., Kloosterman, W. P., Rowland, B. D., de Wit, E., de Ridder, J. & de Laat, W. Enhancer hubs and loop collisions identified from single-allele topologies. Nat. Genet. 50, 1151–1160 (2018).

24. Wu, H., Zhang, J., Tan, L. & Xie, X. S. Single-cell Micro-C profiles 3D genome structures at high resolution and characterizes multi-enhancer hubs. Nat. Genet. 57, 1777–1786 (2025).

25. Montavon, T., Soshnikova, N., Mascrez, B., Joye, E., Thevenet, L., Splinter, E., de Laat, W., Spitz, F. & Duboule, D. A Regulatory Archipelago Controls Hox Genes Transcription in Digits. Cell 147, 1132–1145 (2011).

26. Madsen, J. G. S., Madsen, M. S., Rauch, A., Traynor, S., Van Hauwaert, E. L., Haakonsson, A. K., Javierre, B. M., Hyldahl, M., Fraser, P. & Mandrup, S. Highly interconnected enhancer communities control lineage-determining genes in human mesenchymal stem cells. Nat. Genet. 52, 1227–1238 (2020).

27. Hnisz, D., Shrinivas, K., Young, R. A., Chakraborty, A. K. & Sharp, P. A. A Phase Separation Model for Transcriptional Control. Cell 169, 13–23 (2017).

28. Cho, W.-K., Spille, J.-H., Hecht, M., Lee, C., Li, C., Grube, V. & Cisse, I. I. Mediator and RNA polymerase II clusters associate in transcription-dependent condensates. Science (1979). 361, 412– 415 (2018).

29. Sabari, B. R., Dall’Agnese, A., Boija, A., Klein, I. A., Coffey, E. L., Shrinivas, K., Abraham, B. J., Hannett, N. M., Zamudio, A. V., Manteiga, J. C., Li, C. H., Guo, Y. E., Day, D. S., Schuijers, J., Vasile, E., Malik, S., Hnisz, D., Lee, T. I., Cisse, I. I., Roeder, R. G., Sharp, P. A., Chakraborty, A. K. & Young, R. A. Coactivator condensation at super-enhancers links phase separation and gene control. Science (1979). 361, eaar3958 (2018).

30. Du, M., Stitzinger, S. H., Spille, J.-H., Cho, W.-K., Lee, C., Hijaz, M., Quintana, A. & Cissé, I. I. Direct observation of a condensate effect on super-enhancer controlled gene bursting. Cell 187, 331–344.e17 (2024).

31. de Laat, W. & Duboule, D. Topology of mammalian developmental enhancers and their regulatory landscapes. Nature 502, 499–506 (2013).

32. Bonev, B., Mendelson Cohen, N., Szabo, Q., Fritsch, L., Papadopoulos, G. L., Lubling, Y., Xu, X., Lv, X., Hugnot, J. P., Tanay, A. & Cavalli, G. Multiscale 3D Genome Rewiring during Mouse Neural Development. Cell 171, 557–572.e24 (2017).

33. Caputo, L., Witzel, H. R., Kolovos, P., Cheedipudi, S., Looso, M., Mylona, A., Van Ijcken, W. F. J., Laugwitz, K. L., Evans, S. M., Braun, T., Soler, E., Grosveld, F. & Dobreva, G. The Isl1/Ldb1 Complex Orchestrates Genome-wide Chromatin Organization to Instruct Differentiation of Multipotent Cardiac Progenitors. Cell Stem Cell 17, 287–299 (2015).

34. Oudelaar, A. M., Beagrie, R. A., Gosden, M., de Ornellas, S., Georgiades, E., Kerry, J., Hidalgo, D., Carrelha, J., Shivalingam, A., El-Sagheer, A. H., Telenius, J. M., Brown, T., Buckle, V. J., Socolovsky, M., Higgs, D. R. & Hughes, J. R. Dynamics of the 4D genome during in vivo lineage specification and differentiation. Nature Communications 11, (2020).

35. Drissen, R., Palstra, R.-J., Gillemans, N., Splinter, E., Grosveld, F., Philipsen, S. & de Laat, W. The active spatial organization of the beta-globin locus requires the transcription factor EKLF. Genes Dev. 18, 2485–90 (2004).

36. Vakoc, C. R., Letting, D. L., Gheldof, N., Sawado, T., Bender, M. A., Groudine, M., Weiss, M. J., Dekker, J. & Blobel, G. A. Proximity among Distant Regulatory Elements at the β-Globin Locus Requires GATA-1 and FOG-1. Mol. Cell 17, 453–462 (2005).

37. Phanstiel, D. H., Van Bortle, K., Spacek, D., Hess, G. T., Shamim, M. S., Machol, I., Love, M. I., Aiden, E. L., Bassik, M. C. & Snyder, M. P. Static and Dynamic DNA Loops form AP-1-Bound Activation Hubs during Macrophage Development. Mol. Cell 67, 1037–1048.e6 (2017).

38. Williamson, I., Kane, L., Devenney, P. S., Flyamer, I. M., Anderson, E., Kilanowski, F., Hill, R. E., Bickmore, W. A. & Lettice, L. A. Developmentally regulated Shh expression is robust to TAD perturbations. Development (Cambridge) 146, (2019).

39. Paliou, C., Guckelberger, P., Schöpflin, R., Heinrich, V., Esposito, A., Chiariello, A. M., Bianco, S., Annunziatella, C., Helmuth, J., Haas, S., Jerkovic, I., Brieske, N., Wittler, L., Timmermann, B., Nicodemi, M., Vingron, M., Mundlos, S. & Andrey, G. Preformed chromatin topology assists transcriptional robustness of Shh during limb development. Proc. Natl. Acad. Sci. U. S. A. 116, 12390–12399 (2019).

40. Andrey, G., Schöpflin, R., Jerković, I., Heinrich, V., Ibrahim, D. M., Paliou, C., Hochradel, M., Timmermann, B., Haas, S., Vingron, M. & Mundlos, S. Characterization of hundreds of regulatory landscapes in developing limbs reveals two regimes of chromatin folding. Genome Res. 27, 223– 233 (2017).

41. Pollex, T., Rabinowitz, A., Gambetta, M. C., Marco-Ferreres, R., Viales, R. R., Jankowski, A., Schaub, C. & Furlong, E. E. M. Enhancer–promoter interactions become more instructive in the transition from cell-fate specification to tissue differentiation. Nat. Genet. 56, 686–696 (2024).

42. Chen, Z., Snetkova, V., Bower, G., Jacinto, S., Clock, B., Dizehchi, A., Barozzi, I., Mannion, B. J., Alcaina-Caro, A., Lopez-Rios, J., Dickel, D. E., Visel, A., Pennacchio, L. A. & Kvon, E. Z. Increased enhancer–promoter interactions during developmental enhancer activation in mammals. Nat. Genet. 56, 675–685 (2024).

43. Rubin, A. J., Barajas, B. C., Furlan-Magaril, M., Lopez-Pajares, V., Mumbach, M. R., Howard, I., Kim, D. S., Boxer, L. D., Cairns, J., Spivakov, M., Wingett, S. W., Shi, M., Zhao, Z., Greenleaf, W. J., Kundaje, A., Snyder, M., Chang, H. Y., Fraser, P. & Khavari, P. A. Lineage-specific dynamic and pre-established enhancer–promoter contacts cooperate in terminal differentiation. Nat. Genet. 49, 1522–1528 (2017).

44. Kittelmann, S., Preger-Ben Noon, E., McGregor, A. P. & Frankel, N. A complex gene regulatory architecture underlies the development and evolution of cuticle morphology in Drosophila. Curr. Opin. Genet. Dev. 69, 21–27 (2021).

45. Payre, F., Vincent, A. & Carreno, S. ovo/svb integrates Wingless and DER pathways to control epidermis differentiation. Nature 400, 271–5 (1999).

46. Arif, S., Kittelmann, S. & McGregor, A. P. From shavenbaby to the naked valley: trichome formation as a model for evolutionary developmental biology. Evol. Dev. 17, 120–6 (2015).

47. Delon, I., Chanut-Delalande, H. & Payre, F. The Ovo/Shavenbaby transcription factor specifies actin remodelling during epidermal differentiation in Drosophila. Mech. Dev. 120, 747–58 (2003).

48. Frankel, N., Erezyilmaz, D. F., McGregor, A. P., Wang, S., Payre, F. & Stern, D. L. Morphological evolution caused by many subtle-effect substitutions in regulatory DNA. Nature 474, 598–603 (2011).

49. McGregor, A. P., Orgogozo, V., Delon, I., Zanet, J., Srinivasan, D. G., Payre, F. & Stern, D. L. Morphological evolution through multiple cis-regulatory mutations at a single gene. Nature 448, 587–90 (2007).

50. Crocker, J., Abe, N., Rinaldi, L., McGregor, A. P., Frankel, N., Wang, S., Alsawadi, A., Valenti, P., Plaza, S., Payre, F., Mann, R. S. & Stern, D. L. Low affinity binding site clusters confer hox specificity and regulatory robustness. Cell 160, 191–203 (2015).

51. Preger-Ben Noon, E., Davis, F. P. & Stern, D. L. Evolved Repression Overcomes Enhancer Robustness. Dev. Cell 39, 572–584 (2016).

52. Tsai, A., Alves, M. R. P. & Crocker, J. Multi-enhancer transcriptional hubs confer phenotypic robustness. Elife 8, (2019).

53. Tsai, A., Muthusamy, A. K., Alves, M. R., Lavis, L. D., Singer, R. H., Stern, D. L. & Crocker, J. Nuclear microenvironments modulate transcription from low-affinity enhancers. Elife 6, (2017).

54. Preger-Ben Noon, E., Sabarís, G., Ortiz, D., Sager, J., Liebowitz, A. & Stern, D. L. Comprehensive analysis of a cis-regulatory region reveals pleiotropy in enhancer function. Cell Rep. 22, 3021–3031 (2018).

55. Schwartzman, O., Mukamel, Z., Oded-Elkayam, N., Olivares-Chauvet, P., Lubling, Y., Landan, G., Izraeli, S. & Tanay, A. UMI-4C for quantitative and targeted chromosomal contact profiling. Nat. Methods 13, 685 (2016).

56. Said-Ahmad, A., Shimron, N., Samach, E. F., Naik, S., Roy, S., Frankel, N. & Preger-Ben Noon, E. Distinct mechanisms decommission redundant enhancers to facilitate phenotypic evolution. Sci. Adv. 12, (2026).

57. Bonn, S., Zinzen, R. P., Perez-Gonzalez, A., Riddell, A., Gavin, A.-C. & Furlong, E. E. M. Cell type-specific chromatin immunoprecipitation from multicellular complex samples using BiTS-ChIP. Nat. Protoc. 7, 978–94 (2012).

58. Le, M. T., McGehee, J., Dunipace, L., Rumph, D. & Stathopoulos, A. Inferring chromatin architecture at a single locus through probabilistic in situ DNA localization. Nat. Commun. 17, 1752 (2026).

59. Dolsten, G. A., Cofer, E. M., Bing, X. Y., Brack, B., Curlin, M., Theesfeld, C. L., Troyanskaya, O. G., Levine, M. S. & Pritykin, Y. 3D chromatin structures precede genome activation in Drosophila embryogenesis. Cell Genomics 5, (2025).

60. Tomancak, P., Beaton, A., Weiszmann, R., Kwan, E., Shu, S. Q., Lewis, S. E., Richards, S., Ashburner, M., Hartenstein, V., Celniker, S. E. & Rubin, G. M. Systematic determination of patterns of gene expression during Drosophila embryogenesis. Genome Biol. 3, (2002).

61. Kudron, M., Gevirtzman, L., Victorsen, A., Lear, B. C., Gao, J., Xu, J., Samanta, S., Frink, E., Tran-Pearson, A., Huynh, C., Vafeados, D., Hammonds, A., Fisher, W., Wall, M., Wesseling, G., Hernandez, V., Lin, Z., Kasparian, M., White, K., Allada, R., Gerstein, M., Hillier, L., Celniker, S. E., Reinke, V. & Waterston, R. H. Binding profiles for 961 *Drosophila* and *C. elegans* transcription factors reveal tissue-specific regulatory relationships. Genome Res. 34, 2319–2334 (2024).

62. Kudron, M. M., Victorsen, A., Gevirtzman, L., Hillier, L. W., Fisher, W. W., Vafeados, D., Kirkey, M., Hammonds, A. S., Gersch, J., Ammouri, H., Wall, M. L., Moran, J., Steffen, D., Szynkarek, M., Seabrook-Sturgis, S., Jameel, N., Kadaba, M., Patton, J., Terrell, R., Corson, M., Durham, T. J., Park, S., Samanta, S., Han, M., Xu, J., Yan, K.-K., Celniker, S. E., White, K. P., Ma, L., Gerstein, M., Reinke, V. & Waterston, R. H. The ModERN Resource: Genome-Wide Binding Profiles for Hundreds of Drosophila and Caenorhabditis elegans Transcription Factors. Genetics 208, 937–949 (2018).

63. Cavalheiro, G. R., Girardot, C., Viales, R. R., Pollex, T., Ngoc Cao, T. B., Lacour, P., Feng, S., Rabinowitz, A. & Furlong, E. E. M. CTCF, BEAF-32, and CP190 are not required for the establishment of TADs in early Drosophila embryos but have locus-specific roles. Sci. Adv. 9, (2023).

64. Oudelaar, A. M., Harrold, C. L., Hanssen, L. L. P., Telenius, J. M., Higgs, D. R. & Hughes, J. R. A revised model for promoter competition based on multi-way chromatin interactions at the α-globin locus. Nature Communications 10, (2019).

65. Jenett, A., Rubin, G. M., Ngo, T.-T. B., Shepherd, D., Murphy, C., Dionne, H., Pfeiffer, B. D., Cavallaro, A., Hall, D., Jeter, J., Iyer, N., Fetter, D., Hausenfluck, J. H., Peng, H., Trautman, E. T., Svirskas, R. R., Myers, E. W., Iwinski, Z. R., Aso, Y., DePasquale, G. M., Enos, A., Hulamm, P., Lam, S. C. B., Li, H.-H., Laverty, T. R., Long, F., Qu, L., Murphy, S. D., Rokicki, K., Safford, T., Shaw, K., Simpson, J. H., Sowell, A., Tae, S., Yu, Y. & Zugates, C. T. A GAL4-driver line resource for Drosophila neurobiology. Cell Rep. 2, 991–1001 (2012).

66. Schindelin, J., Arganda-Carreras, I., Frise, E., Kaynig, V., Longair, M., Pietzsch, T., Preibisch, S., Rueden, C., Saalfeld, S., Schmid, B., Tinevez, J. Y., White, D. J., Hartenstein, V., Eliceiri, K., Tomancak, P. & Cardona, A. Fiji: An open-source platform for biological-image analysis. Nature Methods vol. 9 676–682 Preprint at 10.1038/nmeth.2019 (2012).

67. Ramos-Rodríguez, M., Subirana-Granés, M. & Pasquali, L. UMI4Cats: an R package to analyze chromatin contact profiles obtained by UMI-4C. Bioinformatics (2021) doi:10.1093/bioinformatics/btab392.

