## Supplementary Materials for "Multiple trans-regulators shape enhancer-promoter hub organization at a multi-enhancer locus"

Naik *et al.*

**This PDF file includes:**

Figures S1 to S10.
Supplementary methods S1-S2.

**Other Supplementary Materials for this manuscript include the following:**

Tables S1 to S6.

### Supplementary Figures

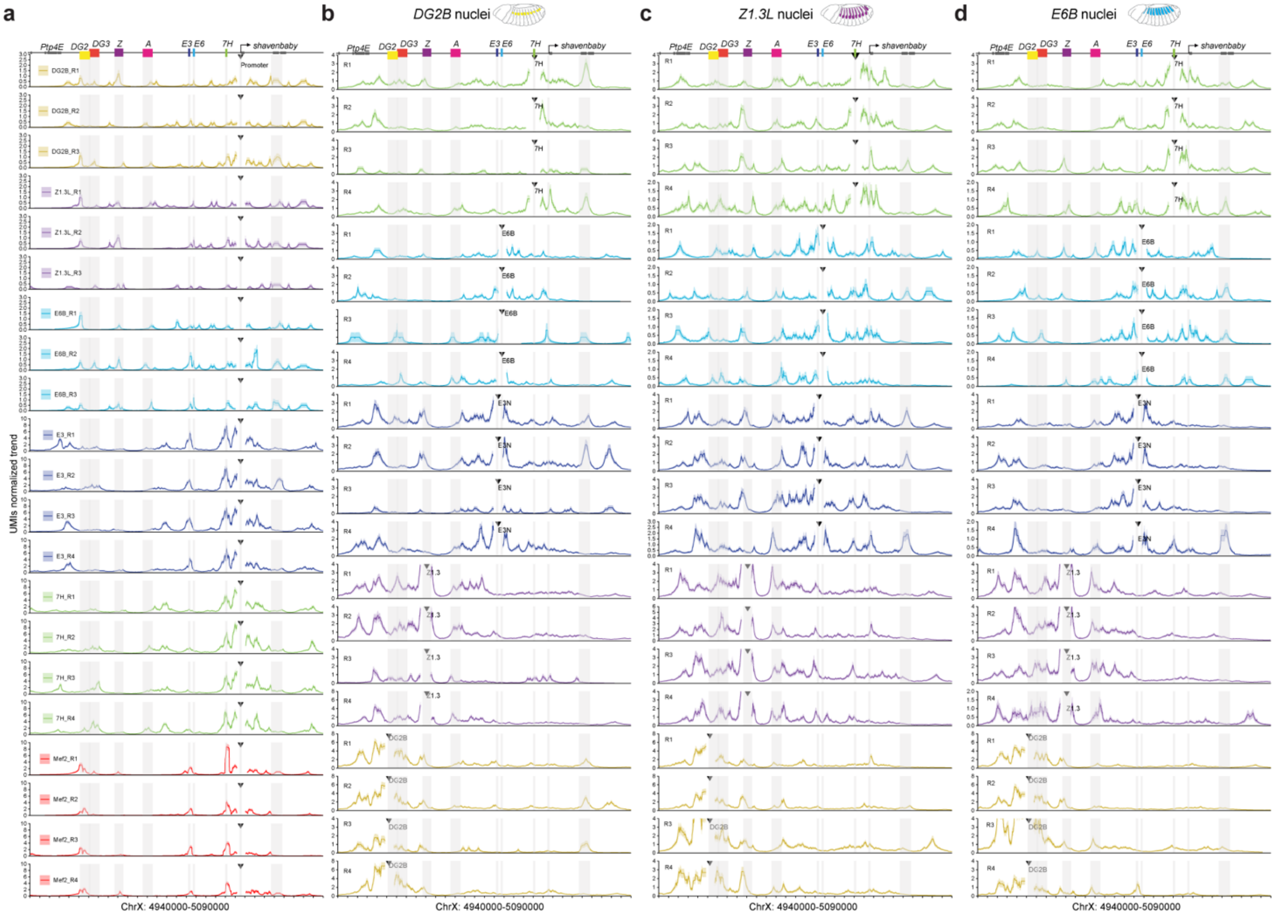


**Figure S1. UMI-4C interaction profiles of biological replicates across the *svb* locus.**

(a) Cell type-specific UMI-4C interaction profiles across the *svb* locus using the endogenous *svb* promoter as the viewpoint. Interaction profiles are shown for *DG2B* (yellow), *Z1.3L* (magenta), *E6B* (cyan), *E3* (blue), *7H* (green), and Mef2 (red) nuclei.

(b–d) Cell type-specific UMI-4C interaction profiles using the indicated *svb* enhancers as viewpoints in *DG2B* (b), *Z1.3L* (c), and *E6B* (d) nuclei. Profiles are shown for the *7H* (green), *E6* (cyan), *E3* (blue), *Z* (magenta), and *DG2* (yellow) enhancer viewpoints.

In all panels, the y-axis shows UMI-normalized interaction frequencies relative to total UMI counts. Gray shading indicates the positions of the *svb* promoter and embryonic enhancers, and triangles indicate the viewpoint used in each interaction profile. Biological replicates are labeled R1-R4.

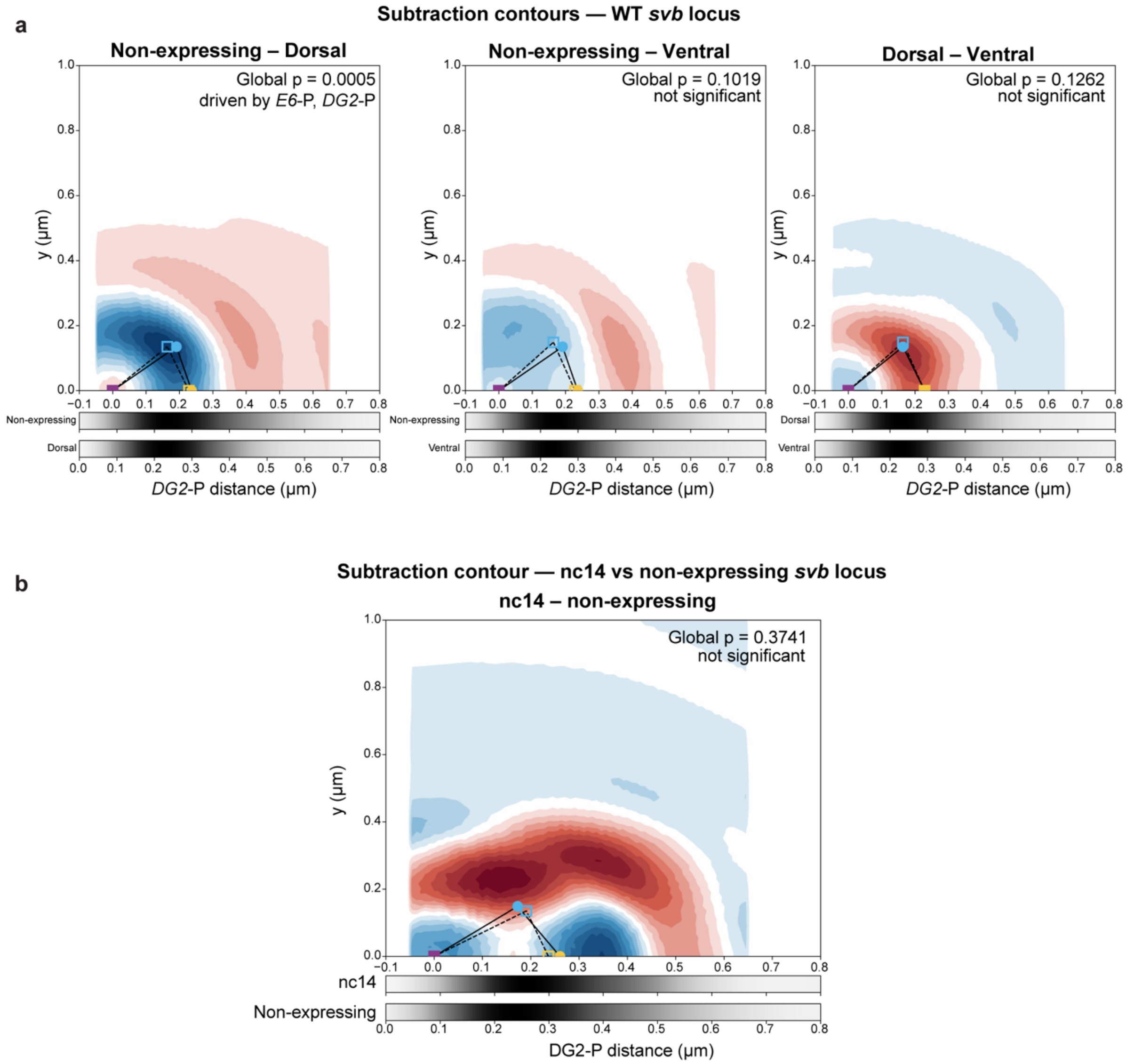


**Figure S2. Probabilistic analysis of three-locus DNA-FISH configurations at the *svb* locus.** Difference contour plots (KDE subtraction) comparing three-locus DNA-FISH configurations between (a) Stage 14 non-expressing and dorsal *svb*-expressing nuclei (left), non-expressing and ventral *svb*-expressing nuclei (middle), and dorsal and ventral *svb*-expressing nuclei (right); and (b) nc14 whole embryos and stage 14 non-expressing nuclei. Red indicates configurations that are more frequent in the first-named population, and blue indicates configurations that are more frequent in the second. Triangles show inferred configurations of the three loci, with the promoter (magenta) set at the origin, *DG2* (yellow) on the x-axis at the modal distance, and *E6* (cyan) at the position of maximum probability. Filled circles connected by solid lines show the first-named population, and open squares connected by dashed lines show the second. Grayscale bars at the bottom show promoter-*DG2* distance distributions. Significance was determined using Simes' method to combine three pairwise KS tests (*E6*-*DG2*, *E6*-promoter, *DG2*-promoter); driving distances are indicated in the top right corner of each panel.

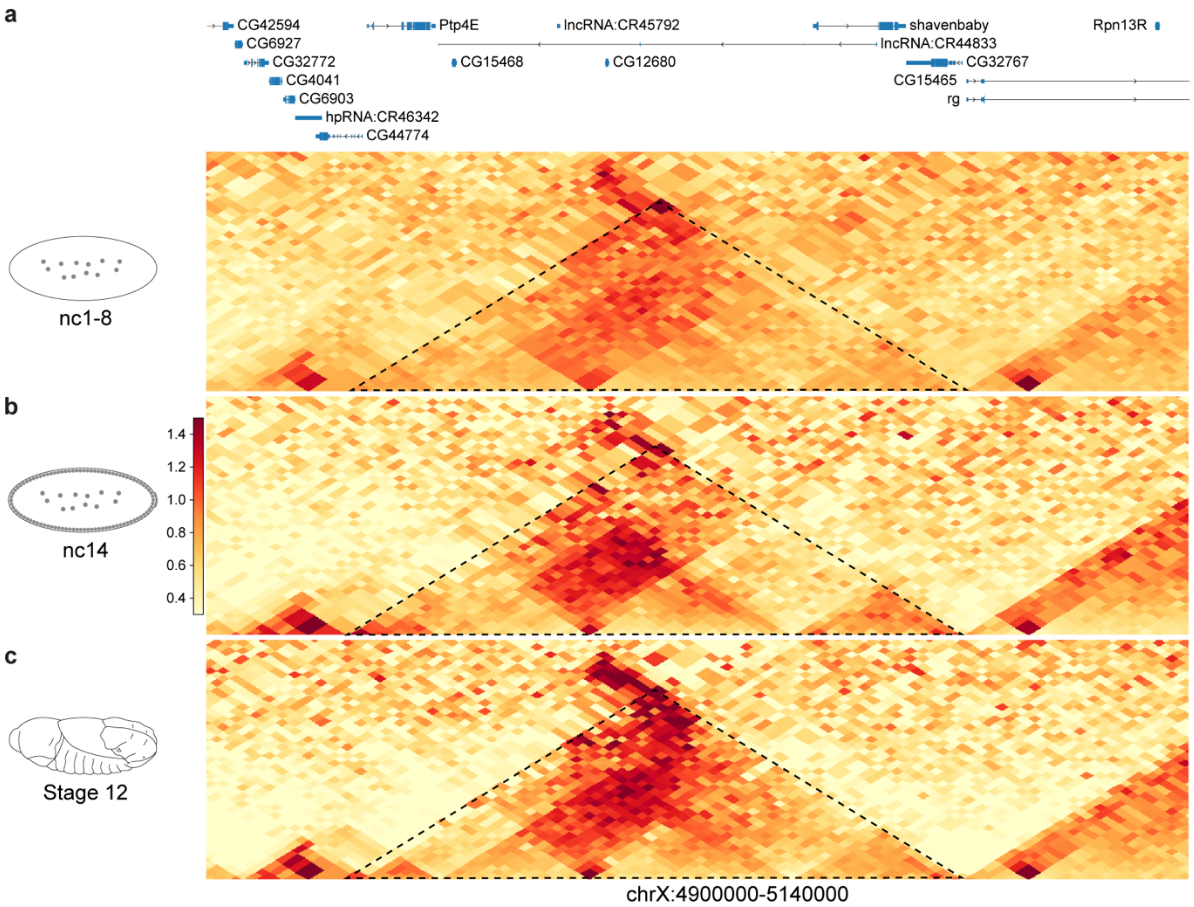


**Figure S3. Chromatin organization at the *svb* locus across embryonic development.** Micro-C contact maps spanning the *svb* locus are shown for embryos at nuclear cycles 1-8 (nc1-8, a), nuclear cycle 14 (nc14, b), and embryonic stages 10-12 (c), using publicly available data from Dolsten et al. 2025[1] (GSE265818). Contact matrices were processed at 10 kb resolution, ICE-normalized, and distance-normalized by observed/expected transformation followed by log1p scaling. At all developmental stages examined, the *svb* locus resides within a well-defined chromatin interaction domain corresponding to the *svb* TAD (indicated by a dashed line).

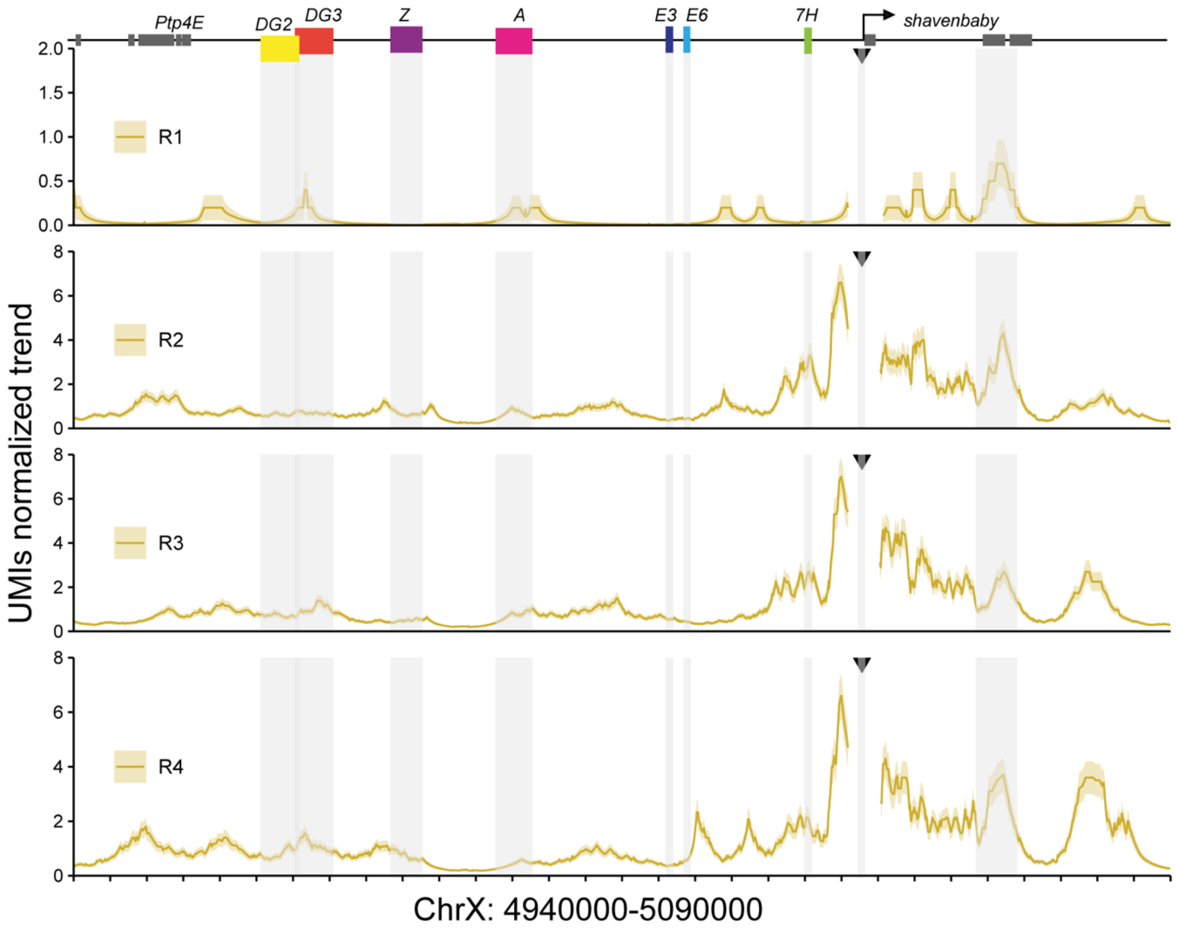


**Figure S4. UMI-4C interaction profiles of biological replicates across the *svb* locus in nc14 embryos.**

UMI-4C interaction profiles generated using the endogenous *svb* promoter as a viewpoint in nuclear cycle 14 (nc14) embryos. The y-axis shows UMI-normalized interaction frequencies relative to the total UMI counts. Gray shading indicates the positions of the *svb* promoter and embryonic enhancers, and triangles indicate the viewpoint used in each interaction profile. Biological replicates are labeled R1-R4.

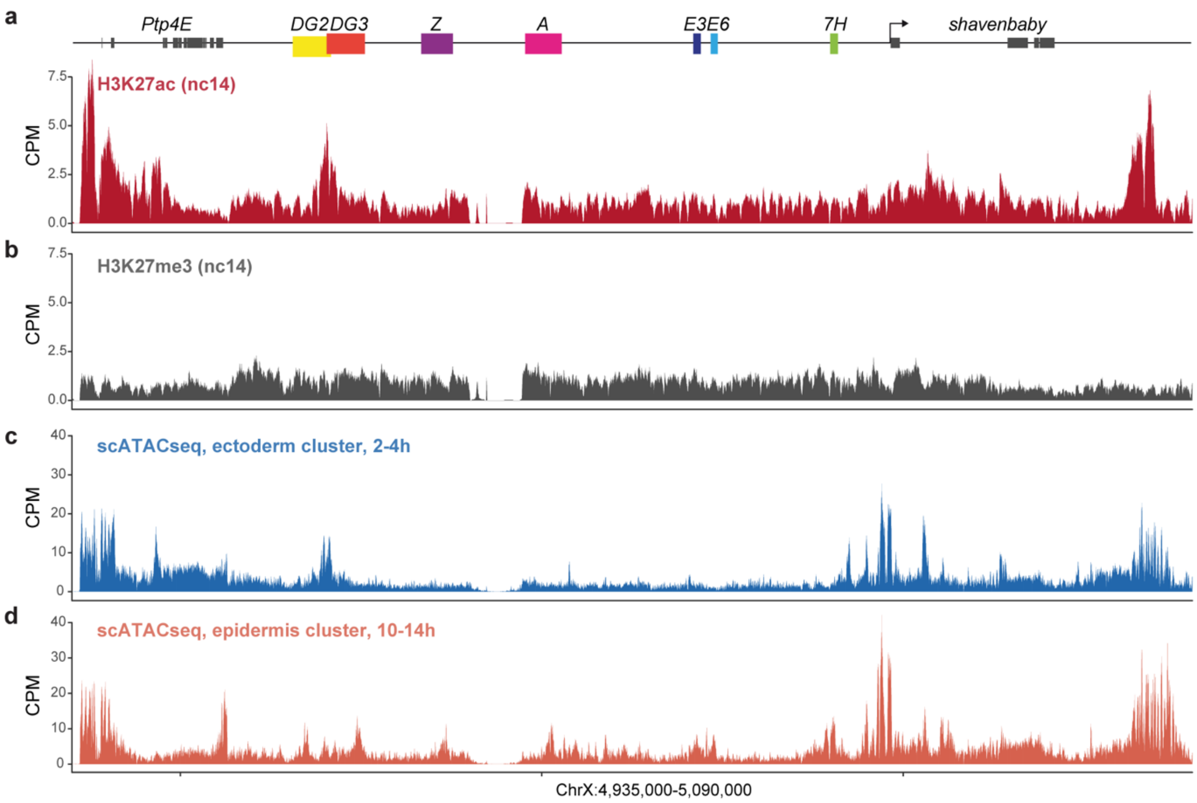


**Figure S5. Histone modification and chromatin accessibility profiles across the *svb* locus.** Genome browser tracks showing chromatin state across the *svb* locus. The positions of the characterized *svb* enhancers are indicated above the tracks.

(a-b) H3K27ac (a) and H3K27me3 (b) ChIP-seq profiles in wild-type nc14 embryos. Data from Gonzaga-Saavedra et al.[2] (GSE299311).

(c-d) CPM-normalized pseudo-bulk scATAC-seq accessibility profile of ectoderm anlage cells (2-4 h, c) and epidermal cells (10-14 h, d). Pseudo-bulk scATAC-seq tracks were generated from the published single-cell ATAC-seq dataset of Calderon et al.[3] (GSE190130) by aggregating accessibility profiles from the indicated cell populations.

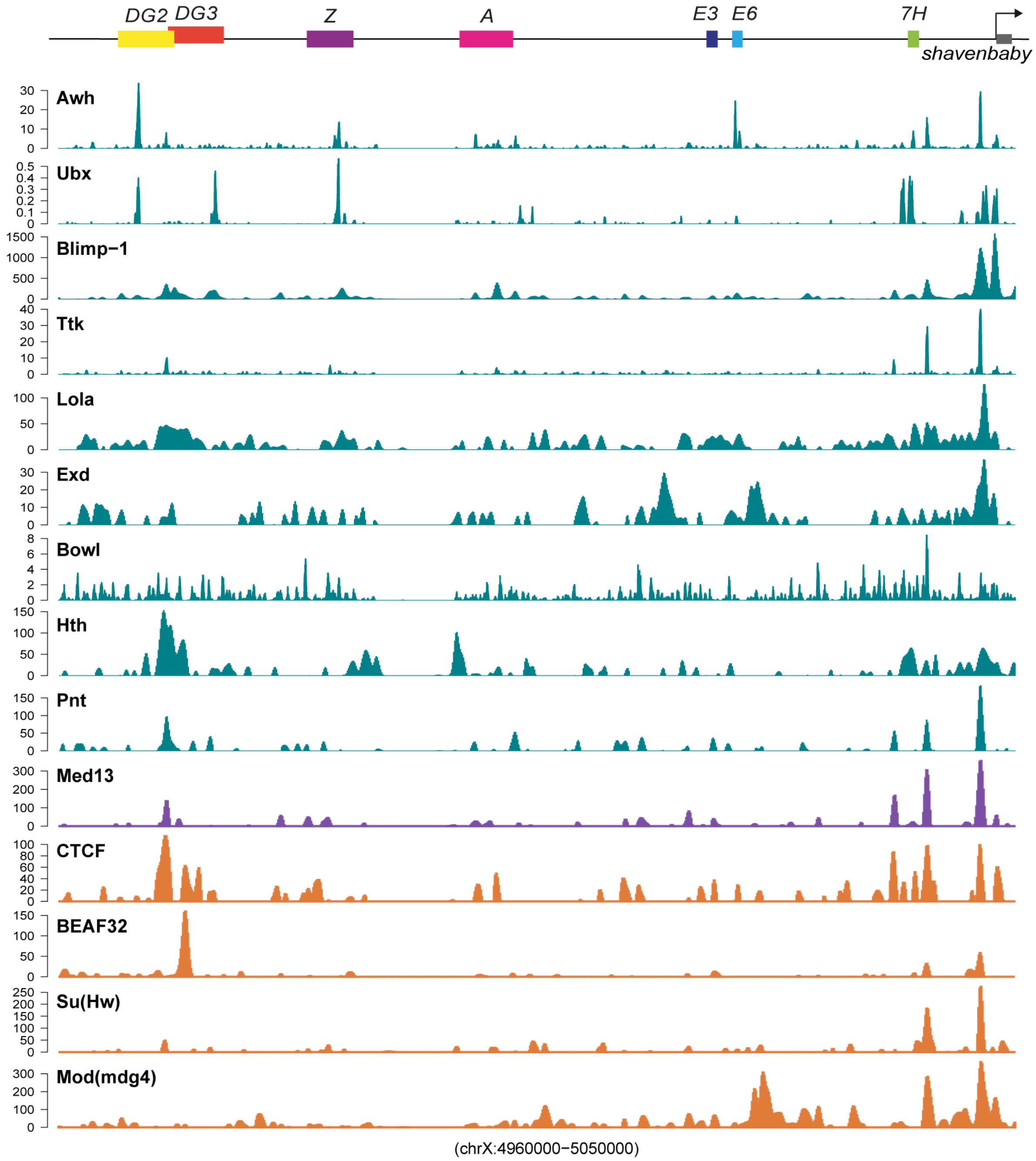


**Figure S6. ChIP-seq analysis of transcription factors and boundary-associated proteins across the *svb* locus.**

Genome browser tracks of ChIP-seq signal for transcription factors (teal), Mediator (purple), and boundary-associated proteins (orange) across the *svb* locus (chrX: 4,960,000-5,050,000). The positions of the characterized *svb* enhancers and the *svb* promoter are indicated above the tracks. All ChIP-seq data were obtained from a study by Kudron et al. [4], except for Ubx, which was obtained from Shlyueva et al. [5](GEO: GSE64284) and reprocessed as described in th Supplementary Methods.

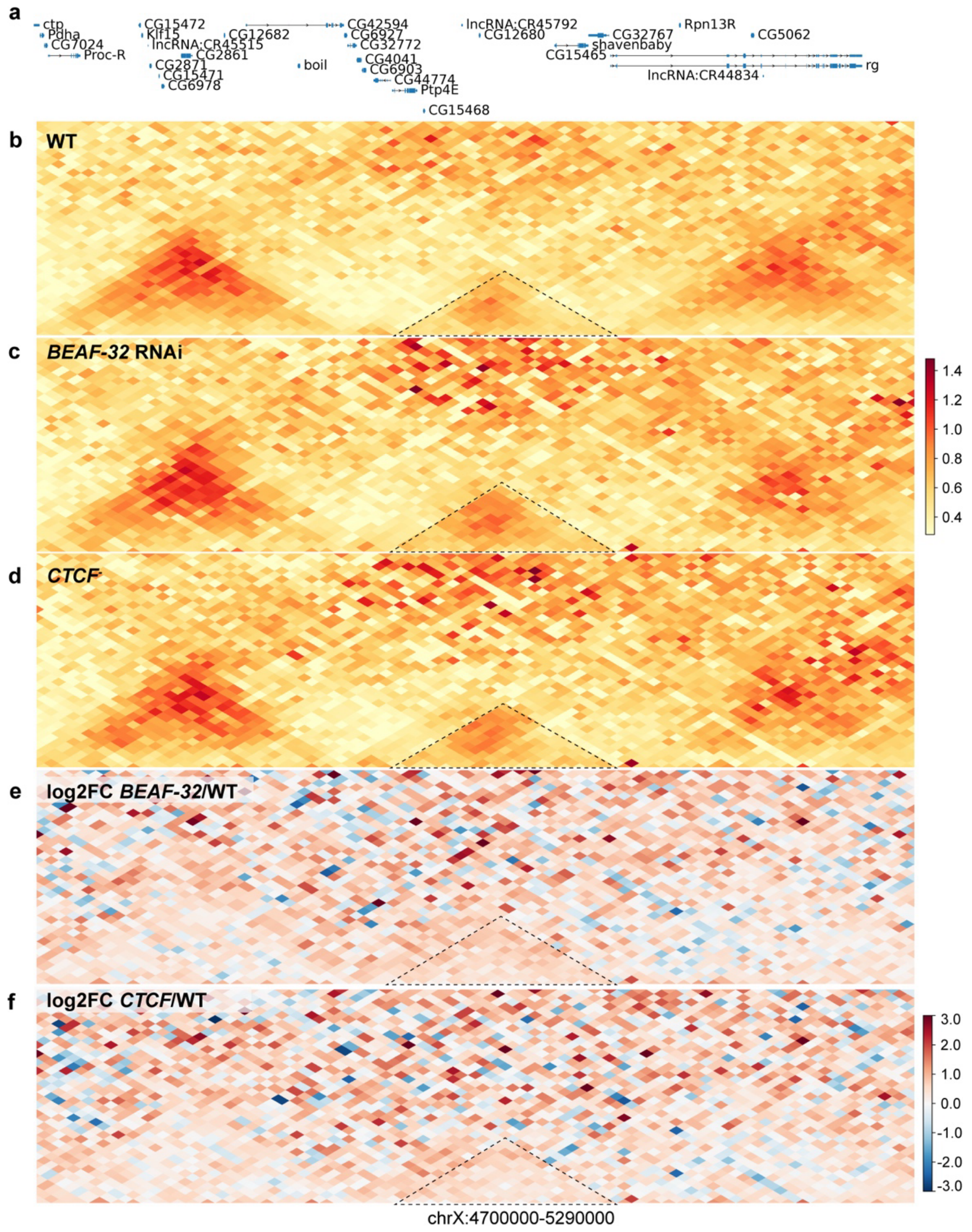


**Figure S7. Depletion of boundary-associated proteins modulate the 3D chromatin organization at the *svb* locus.**

(a) Gene annotation track for the *D. melanogaster* X chromosome region encompassing the *svb* locus (chrX: 4,700,000-5,290,000).

(b-d) Hi-C contact maps (observed/expected, linear scale) for WT embryos (b) and following RNAi-mediated depletion of *BEAF-32* (c), or *CTCF* (d). Warmer colors indicate increased contact enrichment relative to distance-based expectation.

(e-f) $Log_2$ fold-change (RNAi/WT) contact maps for *BEAF-32* (e) and *CTCF* (h), computed from ICE-normalized observed/expected matrices. Differential maps are displayed using a symmetric diverging color scale (red, gain of interactions; blue, loss of interactions), centered at zero. The dashed triangle highlights a subregion within the *svb* locus (chrX: 4,940,000-5,105,000). Hi-C map were generated using data from Cavalheiro et al. [6] (E-MTAB-9158).

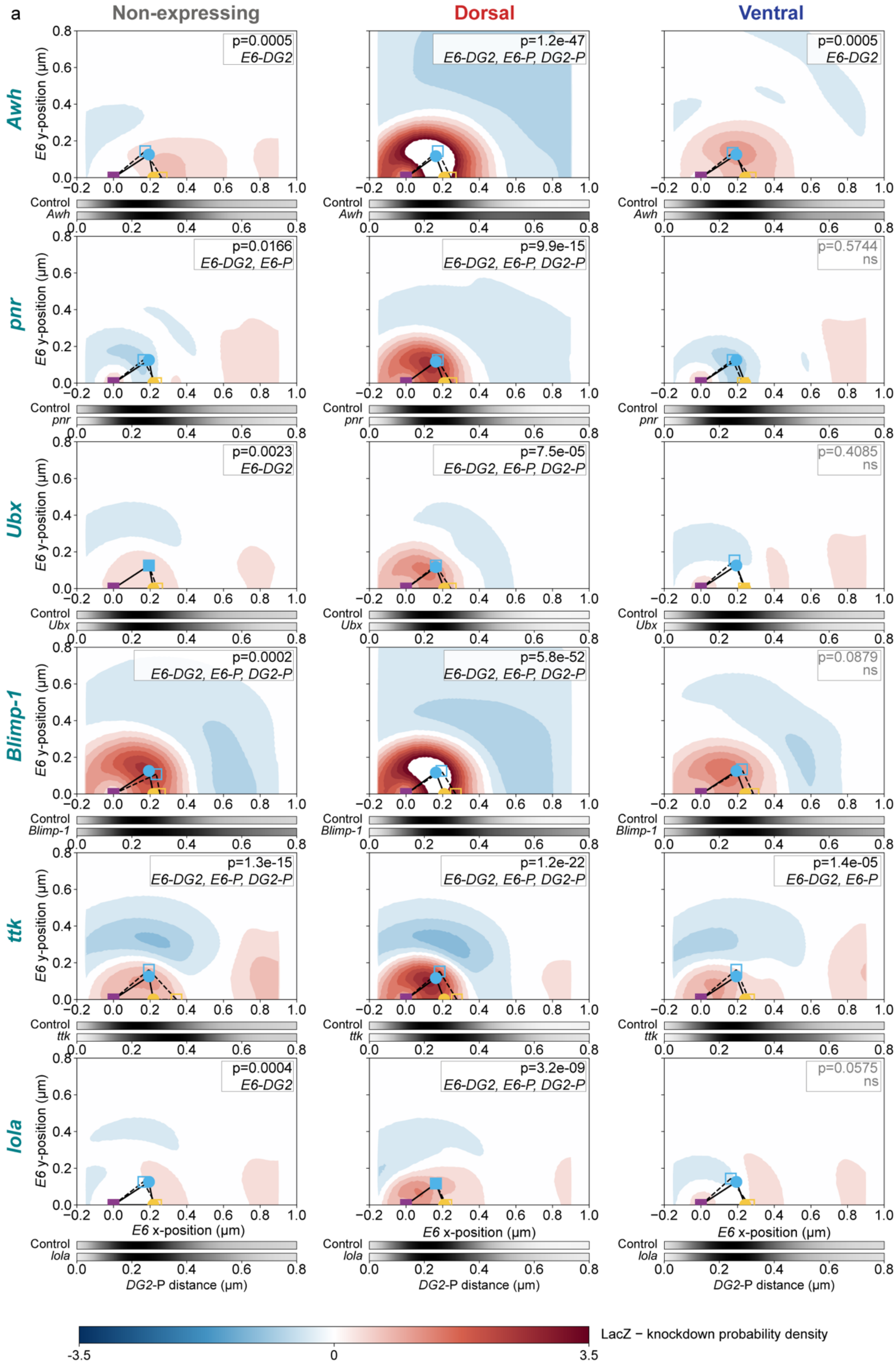

a
Non-expressing
Dorsal
Ventral
Awh
pnr
Ubx
Blimp-1
ttk
lola
E6 y-position (μm)
E6 x-position (μm)
DG2-P distance (μm)
Control
p=0.0005
E6-DG2
p=1.2e-47
E6-DG2, E6-P, DG2-P
p=0.0005
E6-DG2
p=0.0166
E6-DG2, E6-P
p=9.9e-15
E6-DG2, E6-P, DG2-P
p=0.5744
ns
p=0.0023
E6-DG2
p=7.5e-05
E6-DG2, E6-P, DG2-P
p=0.4085
ns
p=0.0002
E6-DG2, E6-P, DG2-P
p=5.8e-52
E6-DG2, E6-P, DG2-P
p=0.0879
ns
p=1.3e-15
E6-DG2, E6-P, DG2-P
p=1.2e-22
E6-DG2, E6-P, DG2-P
p=1.4e-05
E6-DG2, E6-P
p=0.0004
E6-DG2
p=3.2e-09
E6-DG2, E6-P, DG2-P
p=0.0575
ns
LacZ − knockdown probability density
-3.5
0
3.5

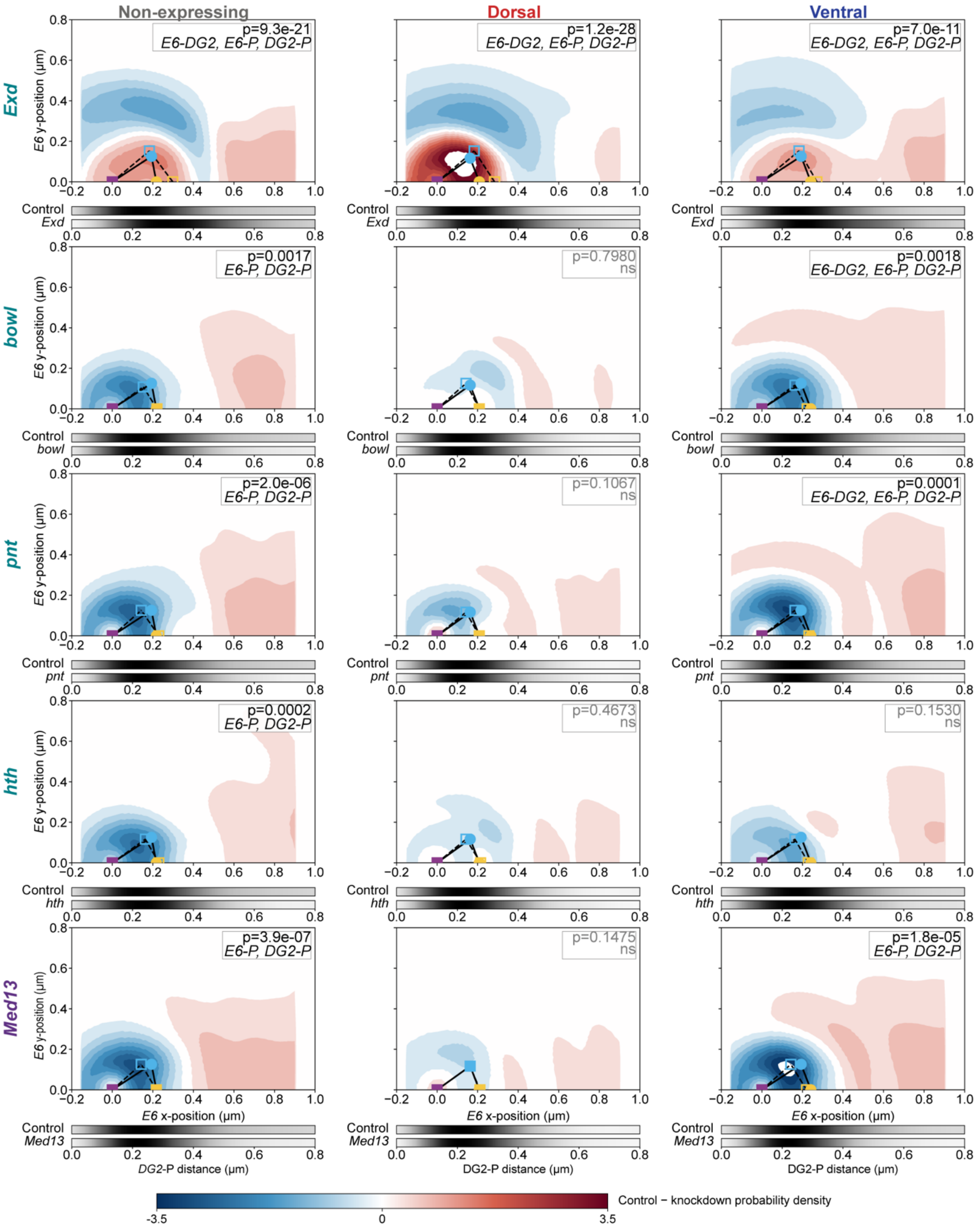

Non-expressing
Dorsal
Ventral
Exd
bowl
pnt
hth
Med13
E6 y-position (µm)
E6 x-position (µm)
DG2-P distance (µm)
Control
p=9.3e-21
E6-DG2, E6-P, DG2-P
p=1.2e-28
E6-DG2, E6-P, DG2-P
p=7.0e-11
E6-DG2, E6-P, DG2-P
p=0.0017
E6-P, DG2-P
p=0.7980
ns
p=0.0018
E6-DG2, E6-P, DG2-P
p=2.0e-06
E6-P, DG2-P
p=0.1067
ns
p=0.0001
E6-DG2, E6-P, DG2-P
p=0.0002
E6-P, DG2-P
p=0.4673
ns
p=0.1530
ns
p=3.9e-07
E6-P, DG2-P
p=0.1475
ns
p=1.8e-05
E6-P, DG2-P
Control − knockdown probability density
-3.5
0
3.5

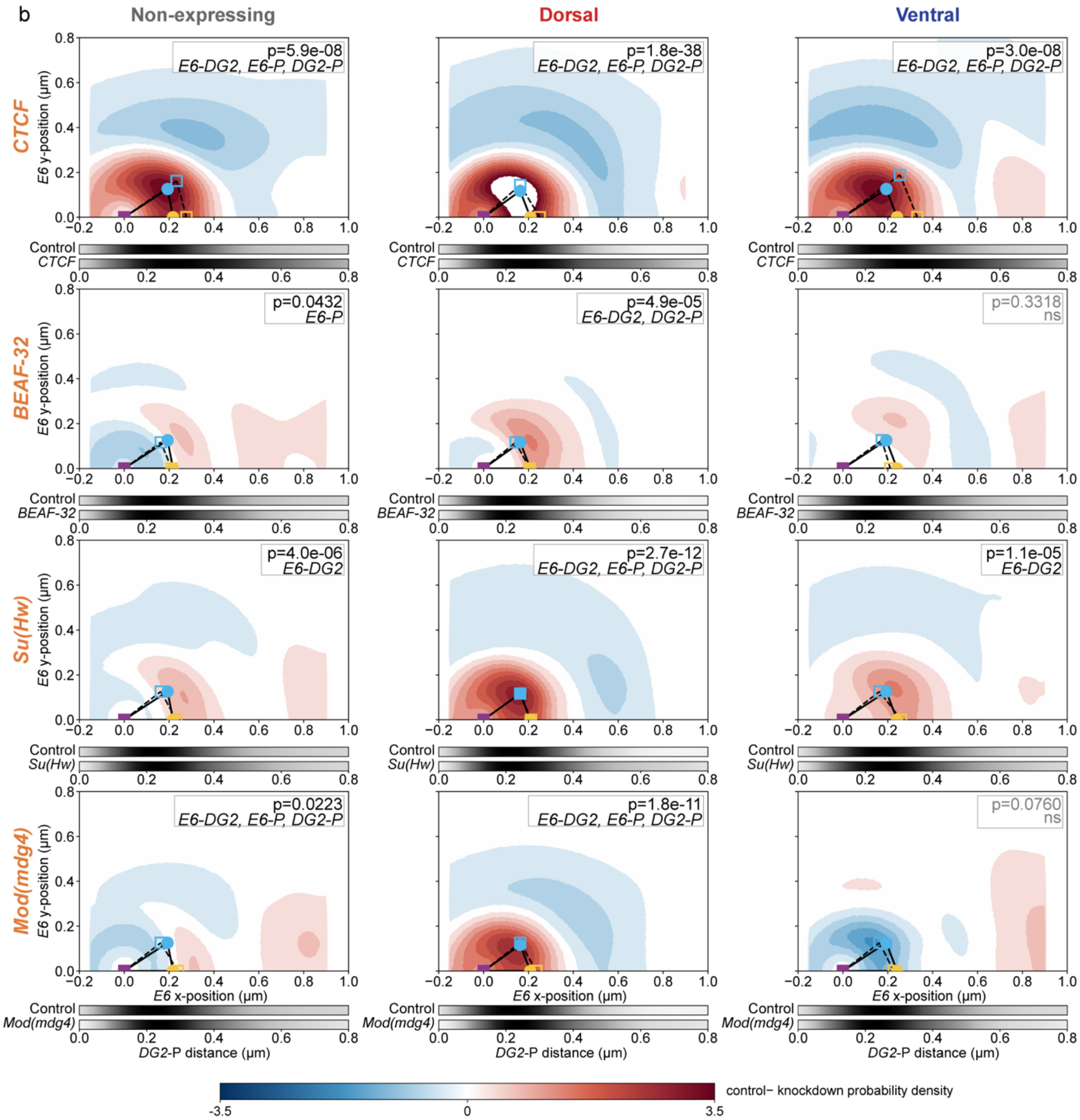


**Figure S8. Probabilistic analysis of three-locus DNA-FISH configurations following RNAi-mediated depletion of candidate *svb* enhancer-promoter hub regulators.** Difference contour plots (KDE subtraction) comparing three-locus DNA-FISH configurations between each RNAi condition and the corresponding *LacZ* control in non-expressing (left), dorsal *svb*-expressing (middle), and ventral *svb*-expressing (right) nuclei. Plots and statistical analyses are as described in Fig. S3, with the *LacZ* control as the first-named population and the RNAi condition as the second.

(a) Transcription factor perturbations: *Awh, Pnr, Ubx, Blimp-1, ttk,* and *lola, Exd, bowl, pnt, hth,* and *Med13.*

(b) Boundary-associated protein perturbations: *CTCF, BEAF-32, Su(Hw),* and *Mod(mdg4)*.

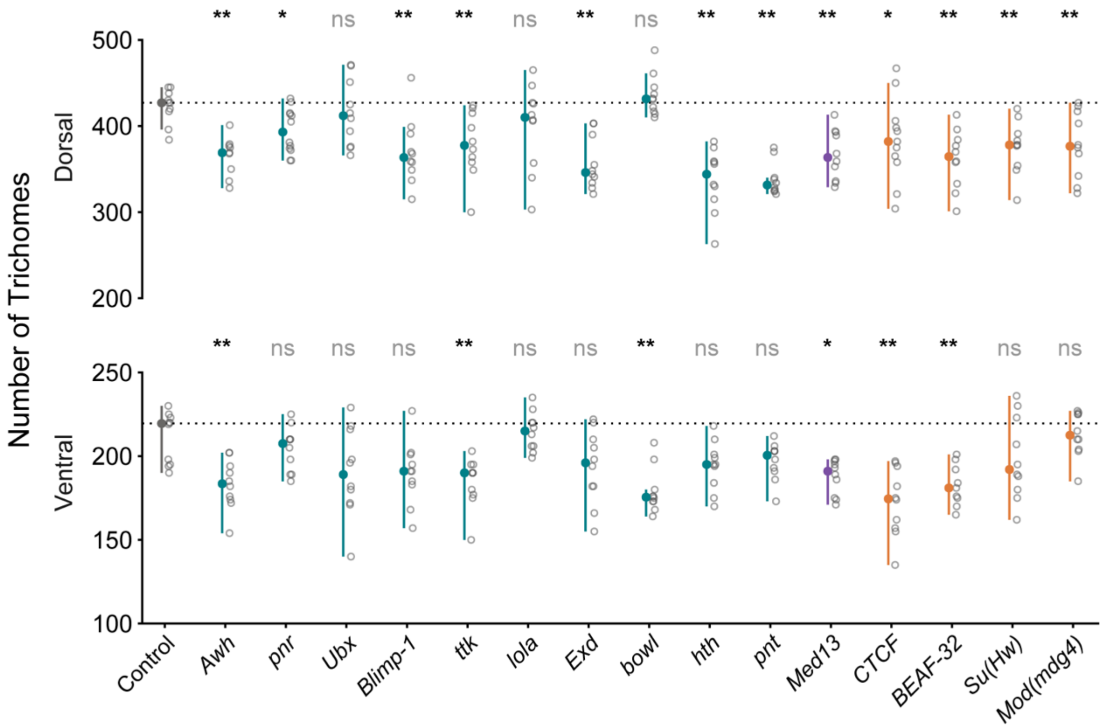


**Figure S9. Trichome numbers following RNAi downregulation of factors contributing to *svb* enhancer-promoter hub organization.**

Number of trichomes on the dorsal (top) and ventral (bottom) cuticle of the fifth abdominal segment of first instar larvae following RNAi-mediated depletion of the indicated transcription factors (teal), Mediator subunit (purple), and boundary-associated proteins (orange), compared with the *LacZ* control (black). Open circles indicate counts from individual larvae, colored circles indicate the mean, and vertical lines denote the standard deviation. Statistical significance was assessed relative to the *LacZ* control using two-sided statistical tests with Benjamini–Hochberg correction. ns, not significant; *****, $P < 0.05$; ******, $P < 0.01$; *******, $P < 0.001$.

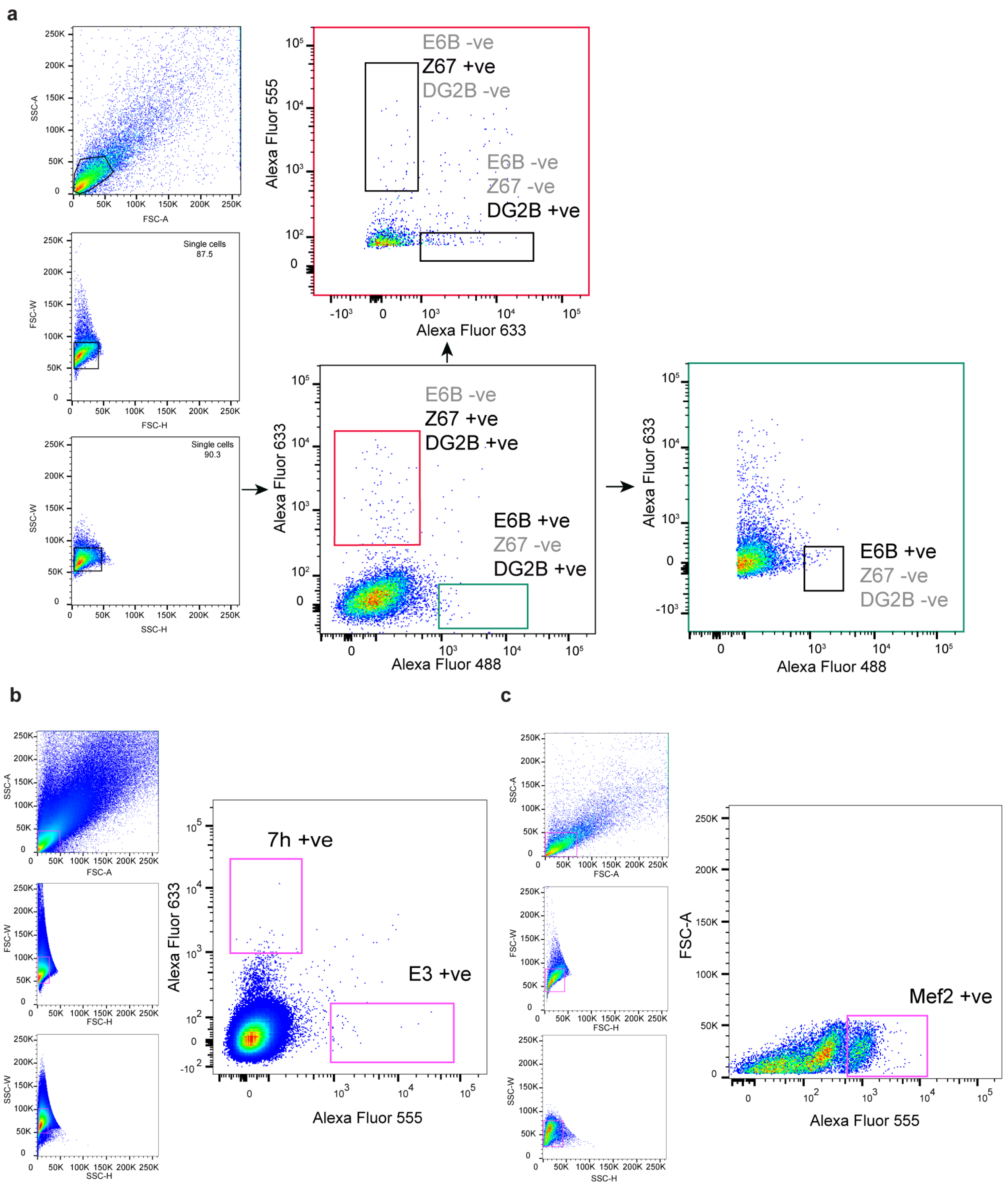


**Figure S10. Fluorescence-activated cell sorting strategy for isolation of specific epidermal subpopulations.**

Nuclei were isolated from cross-linked embryos of the dorsal tester line (a), ventral tester line (c), and the *D. Melanogaster* Oregon$^R$ (b) and stained with antibodies against the corresponding reporter proteins and *Mef2* antibody followed by fluorescently labeled secondary antibodies. For all samples, single nuclei were identified based on forward- (FSC) and side-scatter (SSC) properties followed by doublet exclusion. (a) Sequential gating on Alexa Fluor 488 (*E6B*), Alexa Fluor 555 (*DG2B*), and Alexa Fluor 633 (*Z1.3L*) fluorescence was used to isolate each reporter-positive population while excluding nuclei positive for the other reporter transgenes. (b) Following selection of intact single nuclei and doublet exclusion, *7H*-positive nuclei were identified based on Alexa Fluor 633 fluorescence and *E3*-positive nuclei based on Alexa Fluor 555 fluorescence. (c) *Mef2*-positive nuclei were identified based on Alexa Fluor 555 fluorescence.

### Supplementary methods

#### S1. DNA-FISH data analysis

##### Nuclei segmentation

Nuclei were segmented from the DAPI channel using Cellpose (v4.2.1, default pretrained model, 3D mode; diameter parameter = 30 px, flow threshold = 0.4); to accommodate GPU/memory constraints, each image volume was smoothed (Gaussian, $\sigma = 2$ per z-slice) and downsampled to a fixed 12 × 1024 × 1024 voxel grid prior to segmentation, and the resulting label mask was rescaled to the native image dimensions of each acquisition using nearest-neighbour interpolation to preserve integer nucleus identities. FISH spots were localized using RS-FISH (Fiji/ImageJ v2.9.0; Advanced mode; DoG detection, Gaussian scale $\sigma = 1.8$, anisotropy factor 0.81, RANSAC fitting with maximum error 1.5) applied via a custom batch macro identically across all images. Chromatic shifts between channels were corrected using 0.1 μm TetraSpeck™ microspheres (Thermo Fisher Scientific) imaged under identical optical conditions; a global linear 3D transformation per non-reference channel was calculated by 3D Gaussian bead centroid fitting in the Zeiss ZEN Channel Alignment (Extended) module (633 nm reference) and applied to all experimental datasets prior to spot detection.

##### Three-locus probability density analysis

To visualize and statistically compare the joint spatial configuration of the three loci beyond pairwise distances, two-dimensional probability density surfaces were estimated from the pairwise distance measurements by Gaussian kernel density estimation (KDE), adapting the PLOTTED framework (Le et al. 2026)[7], for each condition or RNAi genotype and the corresponding control. Difference maps were generated by subtracting the KDE surfaces of the two compared populations. The maximum-probability (modal) three-locus configuration for each population was overlaid as a triangle, with the svb promoter fixed at the origin, *DG2* positioned on the x-axis at the modal promoter-*DG2* distance, and *E6* placed at the maximum-probability position inferred from the density surface. Global significance between two populations was calculated using Simes' method to combine three pairwise two-sided Kolmogorov-Smirnov tests (*E6-DG2*, *E6*-promoter, *DG2*-promoter distances).

#### S2. Reanalysis of published datasets

### Hi-C

Published Hi-C data (wild-type, *BEAF-32* RNAi, and *CTCF* depletion; Cavalheiro et al.[6], ArrayExpress: E-MTAB-9158) from early *D. melanogaster* embryos were aligned to dm6 using Juicer (v1.6; BWA v0.7.17, duplicate removal). Contact matrices at 10 kb resolution were ICE-normalized (HiCExplorer v3.7.6), transformed to observed/expected values, and $\log_{1+}$ scaled. Differential maps ($\log_2$[RNAi/WT]) were generated with hicCompareMatrices and visualized with pyGenomeTracks (v3.8) over a 590 kb window spanning the *svb* locus (chrX:4,700,000-5,290,000; dm6).

##### Micro-C

Published Micro-C data spanning early embryogenesis (Dolsten et al.[1], GEO: GSE265818) were used to assess chromatin domain organization at the *svb* locus across nuclear cycles 1-8, nuclear cycle 14, and post-gastrulation stages. Contact matrices at 10 kb resolution were ICE-normalized, distance-normalized by observed/expected transformation, and log1p-scaled prior to visualization.

##### scATAC-seq

Single-cell ATAC-seq data from Calderon et al.[3] (GEO: GSE190130) were reanalyzed to assess chromatin accessibility at the *svb* locus in presumptive ectoderm (2-4 h, ectoderm anlage; n = 25,516 cells) and committed epidermis (10-14 h, cluster 0; n = 15,443 cells). Cell-type annotations were obtained from the dataset's supplementary metadata (atac_meta.rds), and barcodes for each population were extracted using the refined_annotation field and used to subset chromosome X reads from the corresponding deduplicated BAM files (samtools v1.21). CPM-normalized pseudo-bulk bigwig tracks were generated using deepTools bamCoverage (v3.5; 10 bp bins, --normalizeUsing CPM).

##### Histone modification ChIP-seq

H3K27ac (NC14-late; GSM9037186) and H3K27me3 (NC14-mid) ChIP-seq tracks were obtained as CPM-normalized, merged-replicate bigwig files from Gonzaga-Saavedra et al.[2] (GEO: GSE299311).

##### Transcription factor and *Ubx* ChIP-seq

Transcription factor and chromatin-associated protein ChIP-seq binding profiles were obtained from the modERN consortium[4]. Raw *Ubx* ChIP-seq reads (Shlyueva et al.[5], GEO: GSE64284) were aligned to dm6 with Bowtie2, deduplicated, and MAPQ-filtered (≥30). Pooled, input-normalized signal (MACS2, logLR) was converted to bigWig for visualization alongside modERN transcription factor tracks.

All reanalyzed tracks were visualized over the *svb* locus (chrX:4,935,000-5,090,000; dm6) using custom R scripts employing rtracklayer for bigwig import, ggplot2 for coverage rendering, and TxDb.Dmelanogaster.UCSC.dm6.ensGene with org.Dm.eg.db for gene model annotation. Paired tracks (ectoderm anlage versus epidermis; H3K27ac versus H3K27me3) were displayed on a shared y-axis scale to enable direct visual comparison.

#### References:


1. Dolsten, G. A., Cofer, E. M., Bing, X. Y., Brack, B., Curlin, M., Theesfeld, C. L., Troyanskaya, O. G., Levine, M. S. & Pritykin, Y. 3D chromatin structures precede genome activation in Drosophila embryogenesis. *Cell Genomics* **5**, (2025).
2. Gonzaga-Saavedra, N., Degen, E. A., Soluri, I. V, Croslyn, C. & Blythe, S. A. Nucleation-dependent propagation of Polycomb modifications emerges during the Drosophila maternal to zygotic transition. *Elife* **14**, (2025).
3. Calderon, D., Blecher-Gonen, R., Huang, X., Secchia, S., Kentro, J., Daza, R. M., Martin, B., Dulja, A., Schaub, C., Trapnell, C., Larschan, E., O'Connor-Giles, K. M., Furlong, E. E. M. & Shendure, J. The continuum of Drosophila embryonic development at single-cell resolution. *Science (1979).* **377**, (2022).
4. Kudron, M., Gevirtzman, L., Victorsen, A., Lear, B. C., Gao, J., Xu, J., Samanta, S., Frink, E., Tran-Pearson, A., Huynh, C., Vafeados, D., Hammonds, A., Fisher, W., Wall, M., Wesseling, G., Hernandez, V., Lin, Z., Kasparian, M., White, K., Allada, R., Gerstein, M., Hillier, L. D., Celniker, S. E., Reinke, V. & Waterston, R. H. Binding profiles for 961

Drosophila and C. elegans transcription factors reveal tissue-specific regulatory relationships. *Genome Res.* **34**, (2024).

5. Shlyueva, D., Meireles-Filho, A. C. A., Pagani, M. & Stark, A. Genome-wide Ultrabithorax binding analysis reveals highly targeted genomic loci at developmental regulators and a potential connection to polycomb-mediated regulation. *PLoS One* **11**, (2016).
6. Cavalheiro, G. R., Girardot, C., Viales, R. R., Pollex, T., Ngoc Cao, T. B., Lacour, P., Feng, S., Rabinowitz, A. & Furlong, E. E. M. CTCF, BEAF-32, and CP190 are not required for the establishment of TADs in early Drosophila embryos but have locus-specific roles. *Sci. Adv.* **9**, (2023).
7. Le, M. T., McGehee, J., Dunipace, L., Rumph, D. & Stathopoulos, A. Inferring chromatin architecture at a single locus through probabilistic in situ DNA localization. *Nature Communications* **17**, (2026).
